# Limited overlap of eQTLs and GWAS hits due to systematic differences in discovery

**DOI:** 10.1101/2022.05.07.491045

**Authors:** Hakhamanesh Mostafavi, Jeffrey P. Spence, Sahin Naqvi, Jonathan K. Pritchard

## Abstract

Most signals in genome-wide association studies (GWAS) of complex traits point to noncoding genetic variants with putative gene regulatory effects. However, currently identified expression quantitative trait loci (eQTLs) explain only a small fraction of GWAS signals. By analyzing GWAS hits for complex traits in the UK Biobank, and cis-eQTLs from the GTEx consortium, we show that these assays systematically discover different types of genes and variants: eQTLs cluster strongly near transcription start sites, while GWAS hits do not. Genes near GWAS hits are enriched in numerous functional annotations, are under strong selective constraint and have a complex regulatory landscape across different tissue/cell types, while genes near eQTLs are depleted of most functional annotations, show relaxed constraint, and have simpler regulatory landscapes. We describe a model to understand these observations, including how natural selection on complex traits hinders discovery of functionally-relevant eQTLs. Our results imply that GWAS and eQTL studies are systematically biased toward different types of variants, and support the use of complementary functional approaches alongside the next generation of eQTL studies.

## 1 Introduction

Genome-wide association studies (GWAS) have identified thousands of genetic variants linked with a variety of human complex traits and diseases [1]. However, uncovering the functional mechanisms of these GWAS hits remains challenging, notably because ~90% of trait-associated variants lie in noncoding regions [2]. GWAS signals are predominantly located in open chromatin regions in relevant cell types, and are enriched in gene regulatory elements and expression quantitative trait loci (eQTLs). These observations suggest that most GWAS hits are mediated by altering gene regulation of nearby genes [2–7].

Motivated by these observations, many studies have integrated GWAS findings with eQTL mapping to gain functional understanding of trait-associated variants [8–12]. However, despite extensive efforts to catalog eQTLs across diverse sets of biosamples, particularly those conducted by the GTEx consortium [13,14], most GWAS hits are not explained by currently known eQTLs [15–17]. For example, across a number of traits analyzed in the GTEx project, just 43% of GWAS hits and a median of 21% of GWAS hits per trait, colocalized with eQTLs (i.e., indicate the same genetic variant) [14]. Similarly, it has been estimated that only 11% of heritability for complex traits is mediated by gene expression in GTEx tissues [18].

Multiple potential explanations have been proposed for the limited overlap between GWAS hits and eQTLs. One is that some GWAS hits may only be eQTLs in specific contexts, e.g., during development [16, 19–22], in specific cell types [23–26], or in response to physiological stimuli such as immune responses [27–31]. These effects are expected to be absent, or hard to detect, in conventional eQTL assays using post-mortem adult whole tissues. Although hundreds of context-specific eQTLs have been identified, their contribution in explaining trait-associated variants has thus far been modest [17, 31]. For example, eQTLs identified at multiple stages of induced pluripotent stem cell (iPSC) differentiation towards neuronal cell types added ~10% more colocalizations with neurological trait loci beyond GTEx eQTLs [22]. Also in GTEx, cell type-specific eQTLs colocalized with ~8% of GWAS hits [25]. While context-specific effects undoubtedly contribute to the limited colocalization, it remains to be seen how much of the gap can be resolved by deeper sampling of cell types and contexts.

A different type of explanation is that perhaps many trait-relevant eQTLs have not yet been discovered due to incomplete statistical power, even if mapping was performed in the correct contexts [18, 32]. However, others have proposed that eQTL mapping is already “saturated” in well-studied tissues [16].

Alternatively, effects on complex traits could be driven by processes other than gene expression, such as splicing [33] and polyadenylation [34]. However, so far those other mechanisms explain fewer trait-associated variants than do eQTLs [14,31]. Lastly, with current sample sizes, eQTL studies are mainly powered to detect cis-eQTLs (affecting nearby genes), while many trait-relevant variants may act as trans-eQTLs (affecting genes elsewhere in the genome) [35–37]. However, standard models of gene regulation predict that trans-eQTLs should be mediated indirectly through cis effects on nearby genes, and thus such variants should in principle be discoverable as cis-eQTLs [16, 38]. Together, these observations suggest that most GWAS hits are indeed cis-eQTLs, but that many have not yet been discovered in eQTL mapping.

To better understand the lack of overlap between GWAS hits and eQTLs, we analyzed GWAS data for 44 complex traits in the UK Biobank, and eQTL data for 38 tissues in the GTEx data. We show that in fact GWAS hits and eQTLs differ systematically: GWAS hits lie at relatively long distances from transcription start sites (TSSs); they are enriched near genes with a wide variety of functional annotations, such as transcription factors; they are under strong selective constraint; and they typically have complex regulatory landscapes across different tissues and cell types. In contrast, eQTLs are tightly clustered near the TSSs of genes that are typically depleted of most functional annotations, show relaxed selective constraint, and have simpler regulatory landscapes.

We close with a model of variant discovery in GWAS and eQTL assays to explain these observations. We show that even if genetic effects on complex traits were entirely mediated by gene expression, many GWAS hits would not be discovered as significant eQTLs (even in the correct causal contexts). One important reason is that natural selection at constrained genes has very different impacts on GWAS discovery than in eQTL discovery.

In summary, GWAS and eQTL mapping tend to maximize power for different types of variants, and current eQTL mapping has limited discovery at the most functionally-important variants. While further context-specific eQTL studies will help somewhat in explaining GWAS hits, we argue here that these efforts should be complemented by a variety of other functional approaches.

## 2 Results

### Analysis overview

For GWAS analyses, we used publicly available summary statistics for 44 traits in the UK Biobank (Table S1). For eQTL analyses, we used the GTEx v8 data for 38 tissues [14] (Table S1), focusing on cis-eQTLs linked to 18,332 protein-coding genes (Table S2). To make the GWAS hits and eQTLs more comparable, we used identical quality control and SNP selection pipelines for both datasets. We removed lead SNPs in strong LD with protein-altering variants, in order to focus on variants that most likely act through gene expression. This pipeline resulted in 22,119 GWAS hits across traits, and 118,996 eQTLs across all gene-tissue pairs (Tables S3,S4). See Methods for all details.

For each GWAS hit and each eQTL, we evaluated properties of the lead SNP with respect to various SNP and genic features. To study genic features, we linked each GWAS hit to the closest TSS among the same 18,332 protein-coding genes. Although we usually do not know the true causal genes for GWAS hits, recent work has shown that using the nearest gene is an extremely useful proxy [39]. In contrast, for eQTLs, we do of course know the relevant SNP-gene pairs. However to make the analyses comparable between GWAS and eQTLs, in most analyses we masked the true eGenes, and instead linked the eQTL SNP to the *nearest* gene in the same way that we did for GWAS hits. We refer to the genes assigned this way as “GWAS genes” and “eQTL genes” in the rest of this paper.

We observed several systematic differences between eQTL and GWAS lead SNPs (Figure S1): specifically, that eQTL SNPs have higher MAF (median 0.24 compared to 0.2), and are in more gene-dense regions (median 11 genes per Mb compared to 8) than GWAS lead SNPs. These differences could confound our comparisons if they covary with features of interest. Therefore, in all our analyses we included control SNPs, for both GWAS hits and eQTLs, that are matched for MAF, LD score and gene density.

Figure 1 provides an overview of our analysis pipeline and main observations. At the end of the paper, we present models to conceptualize these results.

**Figure 1:**
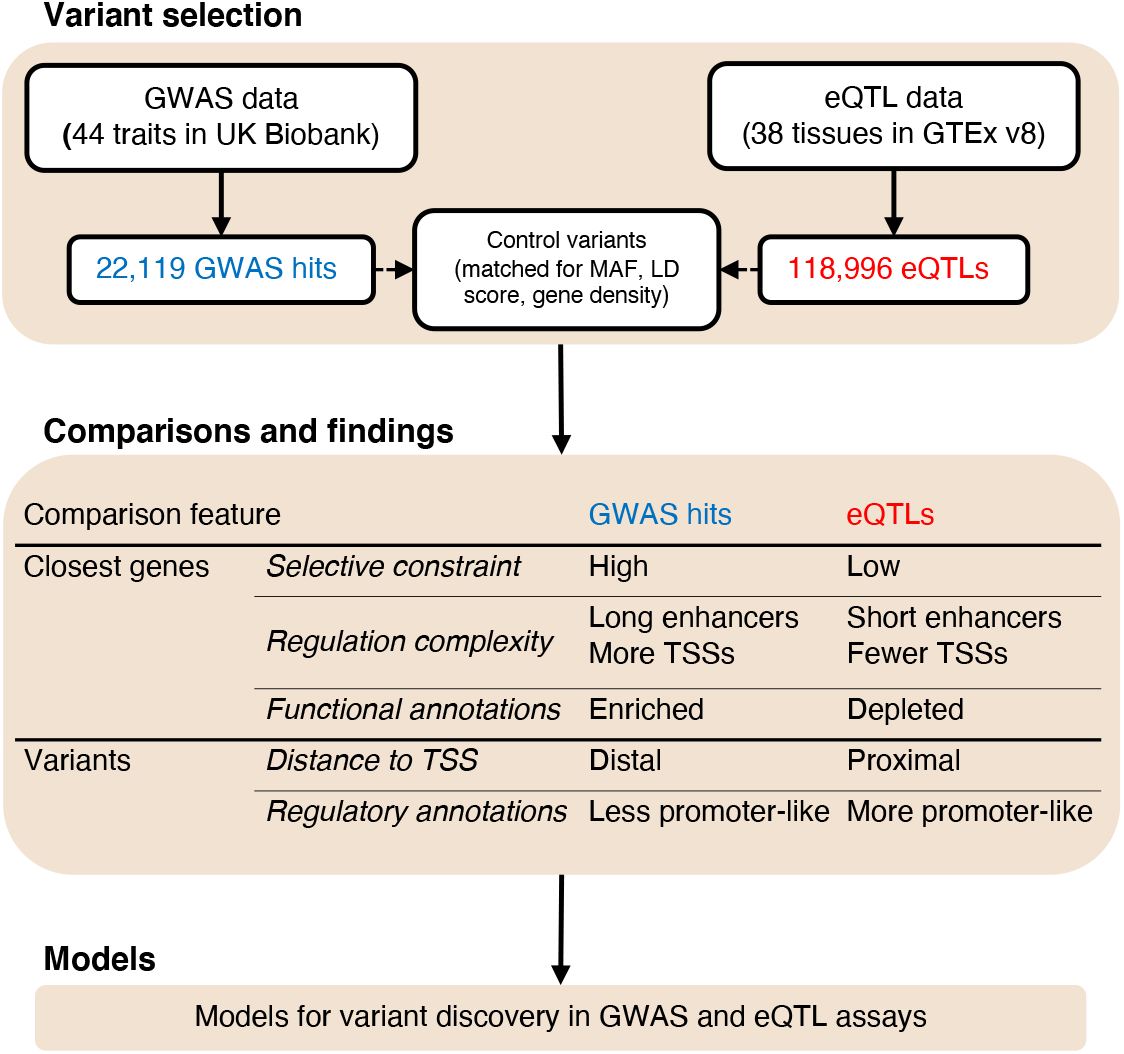
Study workflow and key results. Overview of our pipeline for the analysis of GWAS hits and eQTLs. The list of traits and tissues can be found in Table S1. We compare GWAS hits and eQTLs with respect to a number of genic and SNP features. Summary of our main results is presented in this figure. We describe models for variant discovery in GWAS and eQTL assays to conceptualize these results. See Methods for details.

### Selectively constrained genes are enriched in GWAS genes but depleted in eQTL genes

Previous studies have suggested that the genetic architecture of most complex human traits is shaped by natural selection, such that mutations with large effect sizes are kept at lower frequencies than would be expected in the absence of selection [40–44]. Consistent with selection, SNP heritability for complex traits is enriched near genes depleted of loss-of-function (LoF) variants [45–47], as measured by the pLI score [45]. Although pLI is not an accurate quantitative measure of selection acting on a gene [48], genes with high pLI scores serve as reasonable candidates for being genes under strong selection.

Consistent with previous findings, we observed that GWAS genes are enriched for high-pLI genes (pLI > 0.9) (Figure 2A): 26% of genes linked with GWAS hits compared to 21% for genes linked to matched control SNPs. In contrast, eQTL genes are depleted of high-pLI genes: 12% of genes linked with eQTLs compared to 18% for matched SNPs. These results are in line with previous reports that genes without detectable eQTLs have relatively higher pLI [37, 45].

**Figure 2:**
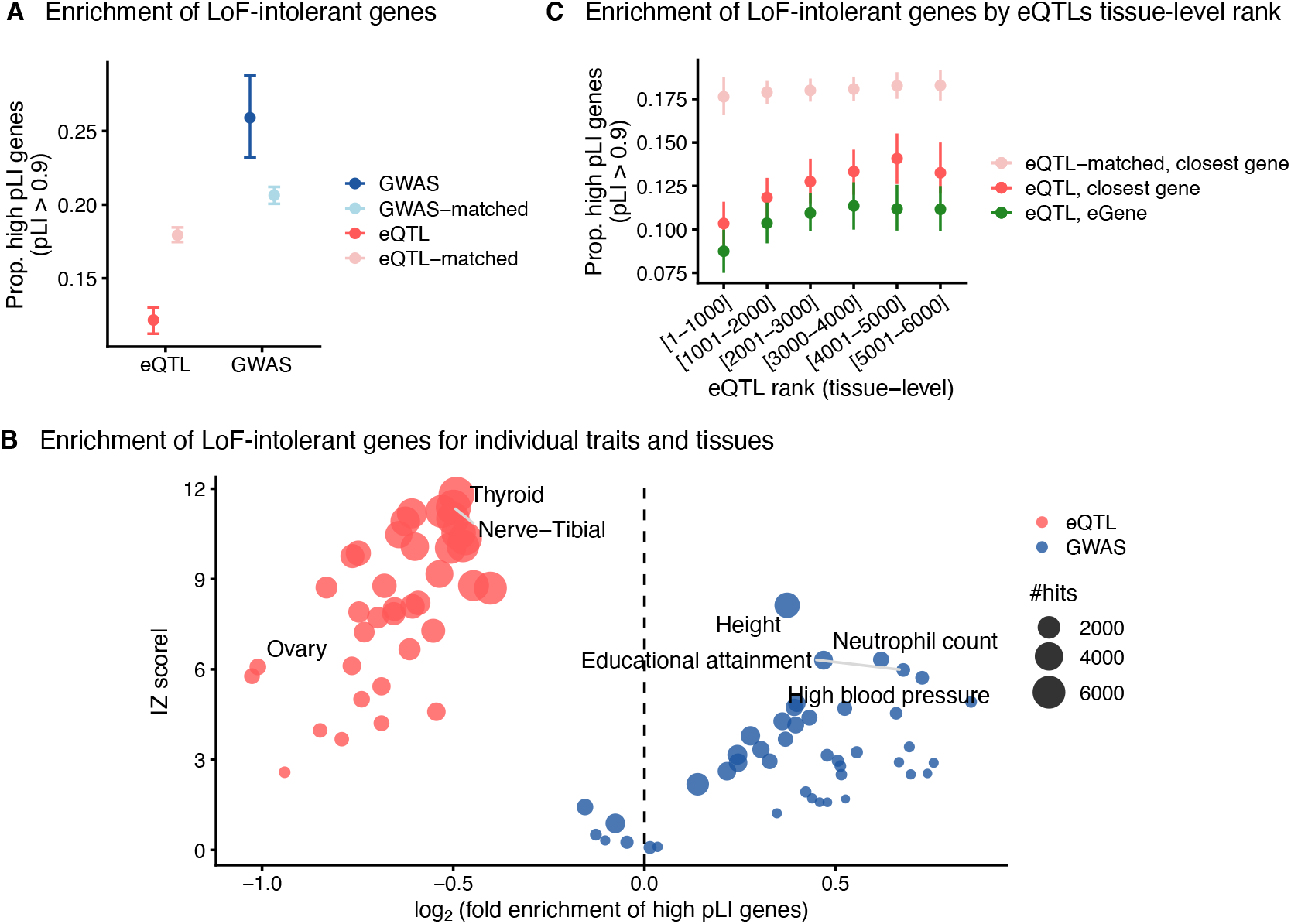
GWAS and eQTL genes are under different selective constraints. **A)** Fraction of genes with high pLI (pLI > 0.9, a measure of selective constraint) among genes nearest to GWAS hits (blue), eQTLs (red), and control SNPs matched for MAF, LD score and gene density. **B)** Enrichment of high-pLI genes in GWAS genes for individual traits and eQTL genes for individual tissues. Enrichment values (on the x-axis) and Z scores (on the y-axis) were computed based on values observed in 1000 sampling iterations of control SNPs. **C)** Fraction of genes with high pLI among eQTL genes (closest gene, red), eGenes (green) and nearest genes to control SNPs (light red) versus eQTL tissue-level rank. For groupings on the x-axis, we first bin ranked eQTLs by association p-values in groups of 1000 eQTLs, and then pooled eQTLs across tissues by the ranked bins. In panels (A) and (C), for GWAS hits and eQTLs, error bars show 95% confidence intervals as determined by quantile bootstrapping. For control SNPs, points and error bars show mean values and 95% confidence intervals in 1000 sampling iterations (see Methods).

In this and most following analyses, we considered GWAS hits pooled across all 44 traits, and eQTLs pooled across all 38 tissues and all genes. We repeated our analysis for each trait/tissue separately, showing that trends in Figure 2A are not driven by a few outlier traits/tissues (Figure 2B). Notably, the depletion of high-pLI genes near eQTLs is replicated in all tissues.

We did, however, see a slight shift towards the null, i.e., less depletion of high-pLI genes, for tissues with larger number of eQTLs (Figure 2B). This is likely due to the inclusion of weaker eQTLs that reach significance detection levels in tissues with larger sample sizes, considering that the depletion of high-pLI genes is increasingly pronounced among top ranked eQTLs (Figure 2C). We observed the same trend for eGenes (true target genes, instead of the closest genes), suggesting that these results are not driven by less accurate gene assignments for weaker eQTLs.

Together, these results show that although a substantial fraction of GWAS hits likely regulate genes that are under strong selection, most discovered eQTLs are not linked with such genes. This suggests that eQTLs with large effects on constrained genes are purged by selection, and is consistent with recent work showing that the fraction of trait heritability estimated to be mediated via gene expression is mostly dominated by genes with low cis-heritability for expression levels [18, 46].

### GWAS genes have more complex regulatory landscapes than eQTL genes

As we showed in Figure 2, genes near GWAS hits and eQTLs differ markedly in terms of their levels of selective constraint. Recently, Wang and Goldstein demonstrated that genes near GWAS hits and eGenes also differ with respect to features of their linked enhancer domains [49]. To explore this further, we considered the transcriptional regulatory landscapes of genes, defined based on the variation in enhancer activity and transcription start site (TSS) usage across diverse tissues and cell types. Many enhancers regulate gene expression in a context-specific manner [50]. Similarly, alternative TSSs could direct context-specific expression, and may also lead to different isoforms [51, 52].

We used the enhancer-gene links in the Roadmap Epigenomics Consortium dataset inferred by Liu et al. [53]. For each gene, we computed (i) the number of active tissue/cell types (out of 111), in which the gene is linked to > 1 enhancer elements, and (ii) the total length of linked enhancers per active tissue/cell type (Figure 3A). We considered these two features jointly using a logistic regression model to distinguish GWAS hits and eQTLs from random SNPs, controlling for MAF, LD score and gene density (Methods). In this framework, compared to genes linked with random SNPs, GWAS genes have longer enhancer regions per active tissue/cell type, while eQTLs have shorter enhancers (Figure 3B), consistent with Wang and Goldstein [49].

**Figure 3:**
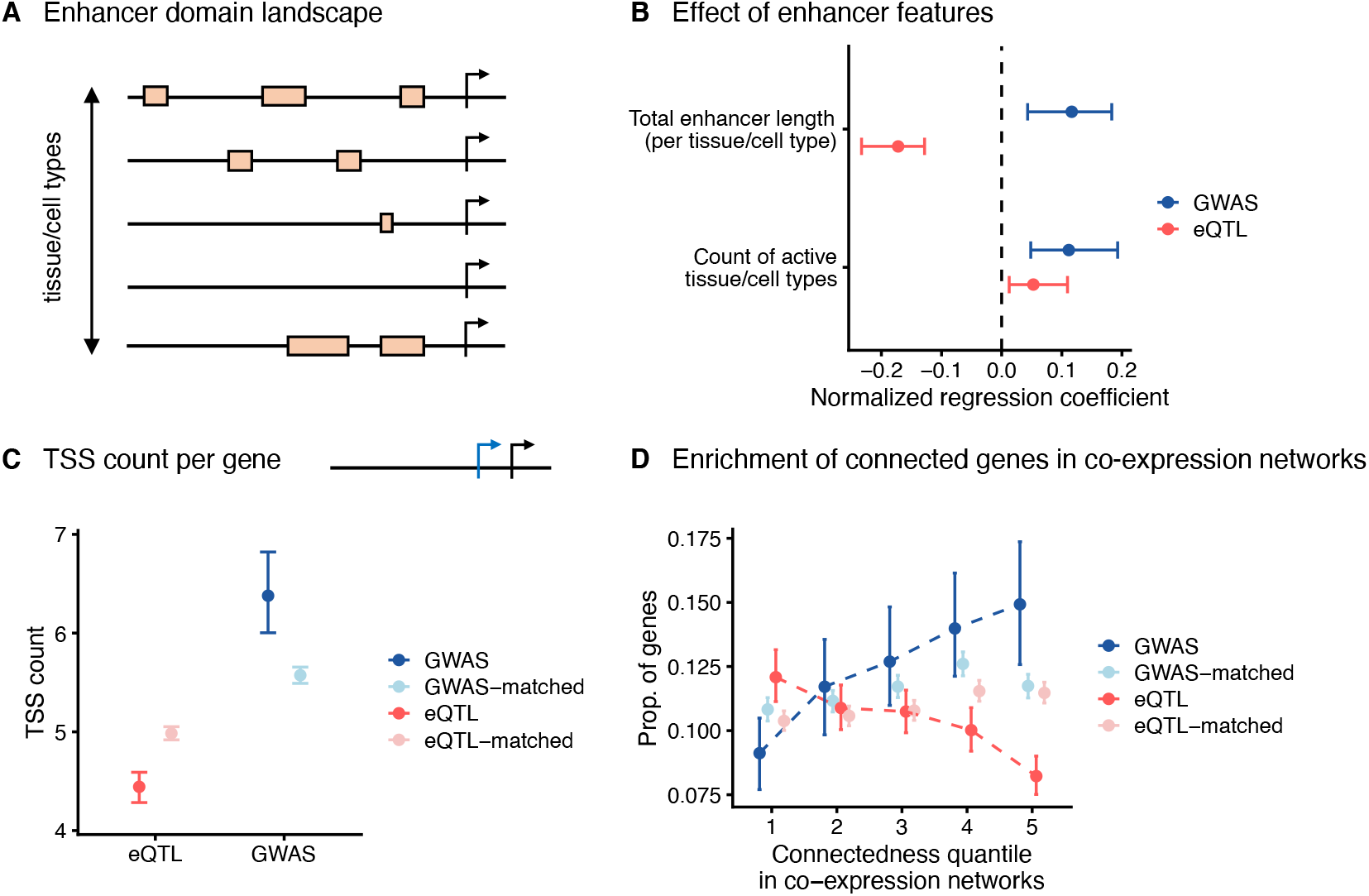
GWAS and eQTL genes have different transcriptional regulatory landscapes. **A)** Schematic of enhancer activity across a number of tissue/cell types. Based on enhancer-gene links inferred from the Roadmap dataset [53], for a given gene, we computed (i) the number of tissue/cell types in which a gene has an enhancer, and (ii) the average total enhancer length (in base pairs) across tissue/cell types with enhancer activity. **B)** Logistic regression coefficients corresponding with the two enhancer features described in (A) for predicting GWAS hits (blue) and eQTLs (red) versus random SNPs after adjusting for confounders (see Methods). **C)** Mean count of TSSs per gene across cell types in the FANTOM project [55] for GWAS and eQTL genes. **D)** Fraction of GWAS and eQTL genes in gene bins ranked by connectedness values computed based on co-expression networks from Saha et al. [56]. In panels (B-D), for GWAS hits and eQTLs, error bars show 95% confidence intervals as determined by quantile bootstrapping. For matched SNPs (for MAF, LD score and gene density, shown in light blue and red colors), points and error bars show mean values and 95% confidence intervals in 1000 sampling iterations.

In contrast, both types of genes have enhancer activity across more tissue/cell types relative to genes linked with random SNPs (Figure 3B), though presumably for unrelated reasons: genes with broad enhancer activity are more likely to harbor regulatory variants in finite sets of tissue/cell types sampled in eQTL datasets. On the other hand, such genes are possibly involved in multiple biological pathways, thus more likely to contribute to complex traits. These results are replicated when using enhancer-gene links from the activity-by-contact model [54] (Figure S2).

We then considered another regulatory feature: how many different TSSs are used for a given gene across a diverse set of cell types. To this end, we computed TSS counts in the FANTOM project [55] (Methods). On average, GWAS genes harbor more TSSs than eQTL genes (6.4 versus 4.4; Figure 3C), and this difference is not explained by the difference observed for matched SNPs (Figure 3C).

We hypothesized that the regulatory landscape of a given gene, in part, corresponds with its regulatory function in a broader gene regulatory network, such that expression of genes with more regulatory domains is more likely to covary with the expression of other genes in the network (e.g., being a target of multiple regulators). Moreover, genes with many downstream regulatory connections in the network, are expected to be important contributors to heritability [35–37]. Motivated by these considerations, we analyzed co-expression networks inferred by Saha et al. for GTEx v6 tissues [56]. For each gene, we constructed a connectedness measure based on the gene’s number of neighbors in tissue-specific networks (Methods). Focusing on 9,422 genes with at least one connection across tissues, we found that genes with more connections are progressively enriched near GWAS hits (Figure 3D), consistent with previous reports that trait heritability is enriched near such genes [57, 58]. In contrast, genes with more connections are progressively depleted for eQTLs (Figure 3D).

Taken together, these observations suggest that targets of GWAS hits are often genes with complex regulatory architecture, in a sense that they harbor regulatory mechanisms to control and diversify gene expression across different contexts and possibly in a context-specific manner, which could correspond with a functional role in gene regulatory networks. These kinds of genes are depleted of eQTLs.

### Many Gene Ontology annotations are enriched among GWAS genes, but depleted among eQTL genes

Differences in the selective constraint and regulatory landscapes of genes likely reflect the functional roles of different types of genes. To gain insight into the role of gene functions in shaping the differences between GWAS and eQTL genes, we analyzed 577 Gene Ontology (GO) biological process terms. For each category, we computed the enrichment of GWAS and eQTL genes in that category compared to control SNPs across all traits and tissues (Methods). For data visualization, we focused here on 41 categories that are broadly unrelated, prioritizing terms that are informative of GWAS/eQTL SNPs (Table S5; Methods).

Notably, we found that GO categories are pervasively enriched among GWAS genes for many traits (Figure 4A). For some categories, the enriched categories highlight evidently relevant traits: e.g., *skeletal system development* for height, *lipid localization* for high cholesterol, and *adaptive immune response* for a number of blood and immune-related traits. That said, many categories, such as *embryonic organ development*, show enrichment across multiple traits.

**Figure 4:**
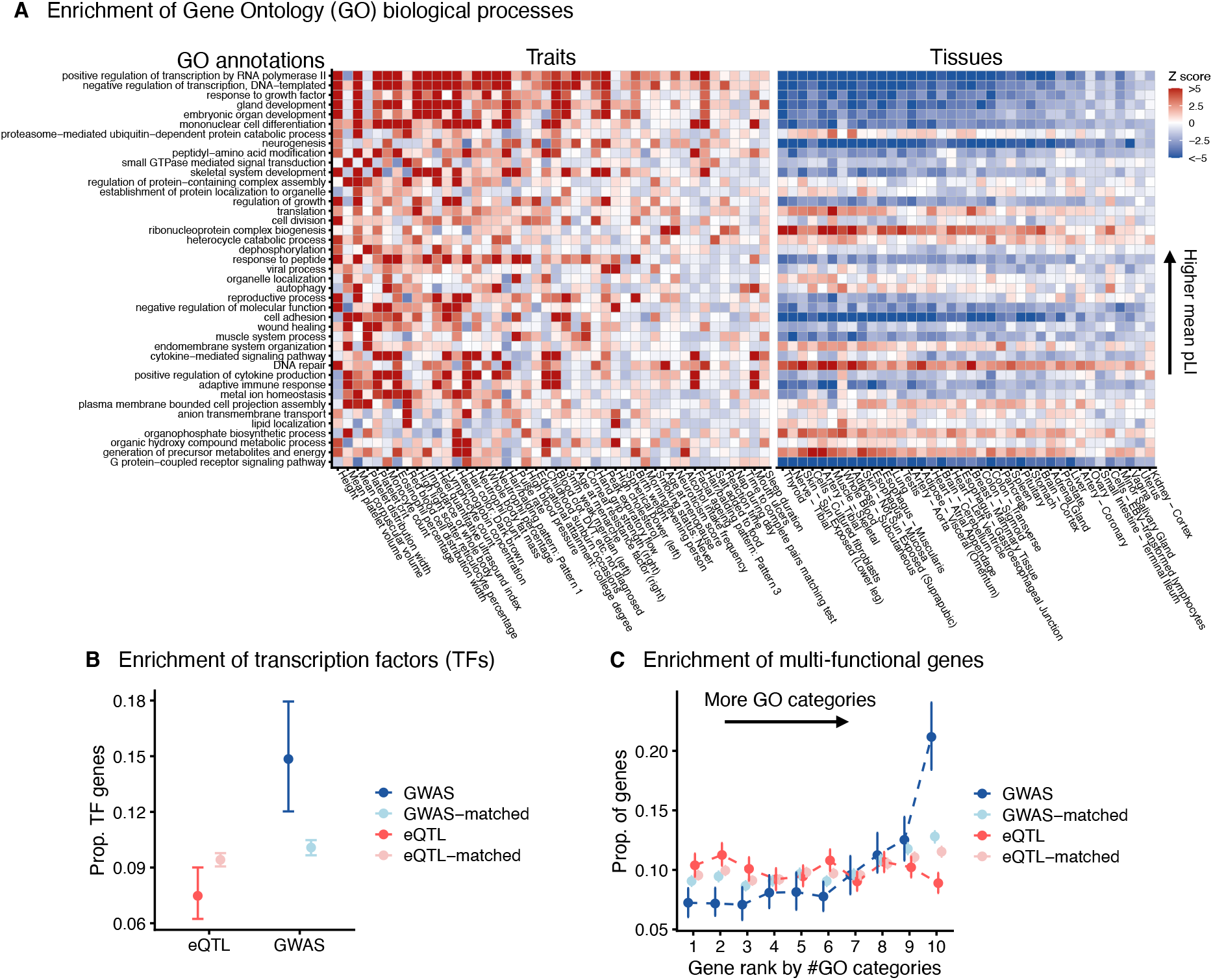
Diverse categories of functional genes are enriched among GWAS genes, but not among eQTL genes. **A)** Enrichment of genes in 41 Gene Ontology (GO) annotations among GWAS and eQTL genes (see Methods for details of GO category selections). Traits and tissues (x-axis) are sorted by hit count (decreasing from left to right), and GO annotations (y-axis) are sorted by the mean pLI value of genes within categories. For each trait- or tissue-GO category pair we computed enrichment Z scores based on 1000 sampling iterations of SNPs matched for MAF, LD score, and gene density (see Methods). The color map represents enrichment (red) or depletion (blue) of a given gene set among GWAS or eQTL genes. See Table S6 for enrichment and Z score values. **B)** Fraction of GWAS and eQTL genes that are transcription factors. **C)** Fraction of GWAS and eQTL genes (y-axis) that are in different bins of genes ranked by the counts of GO categories they belong to (x-axis). In panels (B) and (C), for GWAS hits (blue) and eQTLs (red), error bars show 95% confidence intervals as determined by quantile bootstrapping. For matched SNPs (for MAF, LD score and gene density, shown in light blue and red colors), points and error bars show mean values and 95% confidence intervals in 1000 sampling iterations.

In contrast, many functional categories show clear depletion of eQTLs in all tissues (Figure 4A). Consistent with our earlier results on pLI, depletion is most prominent for categories with high average pLI, suggesting that selection purges eQTLs for genes with evolutionarily important functions. In contrast, certain categories, such as *DNA repair*, are enriched for eQTL genes, suggesting that they may be more tolerant of expression changes.

We wondered whether transcription factors (TFs) would also show the same pattern, as TF dosages often play essential roles in development and cellular functions. Indeed, two categories related to transcriptional regulation, *positive regulation of transcription by RNA polymerase II* and *negative regulation of transcription, DNA-templated*, are particularly enriched in GWAS genes and depleted in eQTL genes (Figure 4A). Focusing specifically on TFs, we found that they are enriched in GWAS genes (15% compared to 10% for matched SNPs), and depleted in eQTL genes (7% compared to 9% for matched SNPs; Figure 4B), in line with previous reports that TFs are underrepresented among eGenes [59]. TFs also contribute substantially to several developmental categories (Figure S3A) that are enriched in GWAS genes and depleted in eQTL genes.

Lastly, we were curious to understand how it is that *most* functional categories can show enrichment for complex traits (Figure 4A). We hypothesized that this is in part because many genes are involved in multiple categories, and such multi-functional genes would be particularly enriched among GWAS genes. Indeed, genes belonging to more GO categories are progressively enriched in GWAS genes, while being modestly depleted for eQTL genes (Figure 4C). Multi-functional genes tend to be highly connected in protein-protein interaction networks (Figure S4A). In line with this observation, we found that genes with the most interactions are enriched for GWAS hits (Figure S4B).

In summary, our GO analysis shows that a diverse range of biological process categories are enriched near GWAS hits; moreover, multi-functional genes are especially enriched for GWAS hits. In contrast, most annotated biological processes are underrepresented near eQTLs.

### GWAS hits are further from transcription start sites (TSSs) than eQTLs

Our analysis so far has focused on differences between GWAS hits and eQTLs with respect to the properties of the target genes. We also found important differences between the SNP-level context of GWAS and eQTL hits.

It’s well known that eQTLs tend to be tightly clustered near TSSs [60–62]. We observed similar results (Figure 5A): 43% of eQTLs lie within 10Kb of the nearest TSS, compared to 23% for matched SNPs. However, in sharp contrast, GWAS hits are only modestly enriched near TSSs (Figure 5A): 22% lie within 10Kb of the nearest TSS, compared to 17% for matched SNPs. GWAS hits typically lie at greater distances from the nearest TSS (median 36Kb) compared to eQTLs (median 13Kb).

**Figure 5:**
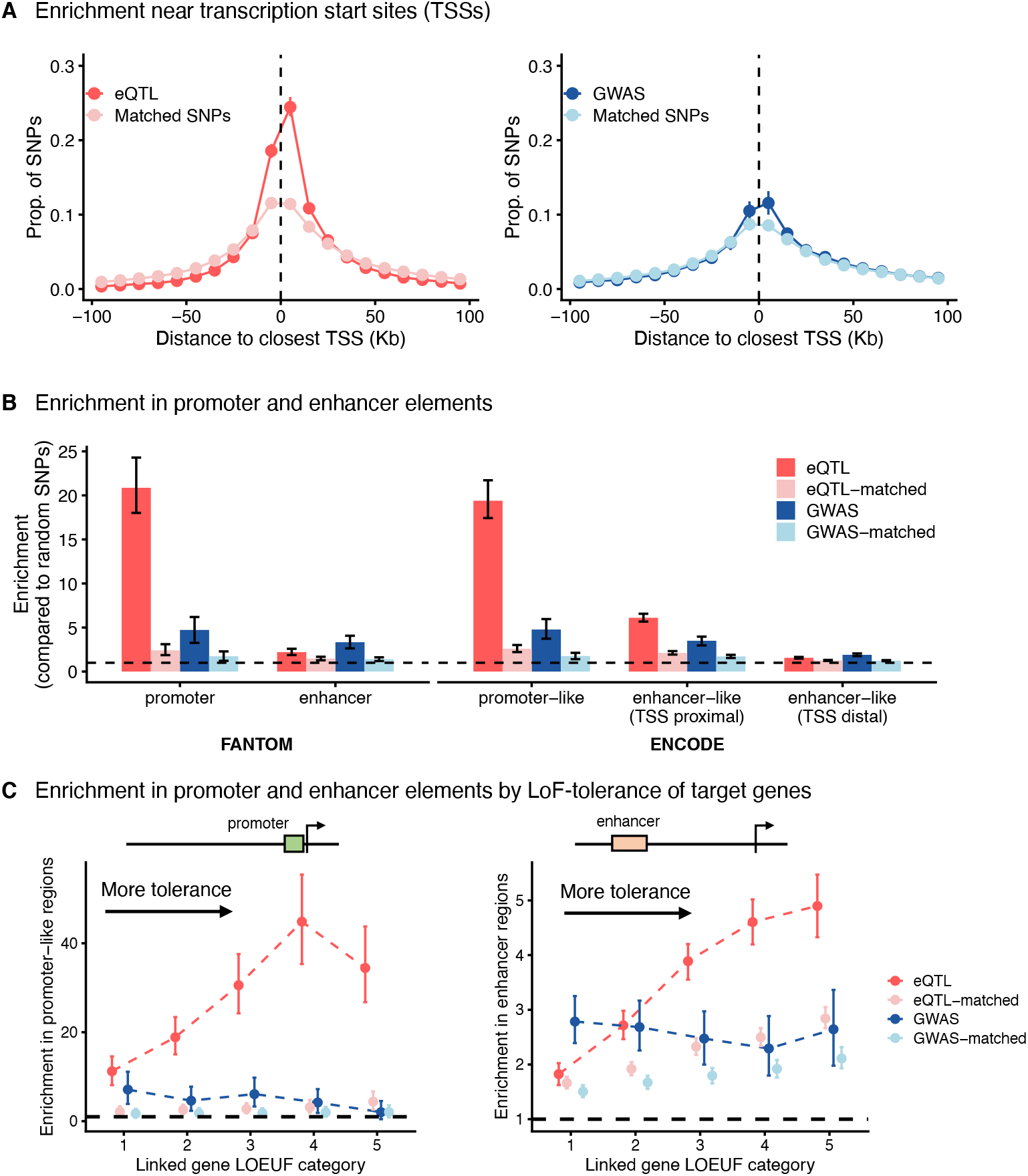
GWAS hits are less enriched at transcription start sites than are eQTLs. **A)** Distance of eQTLs (left panel) and GWAS hits (right panel) to the nearest TSS. Points show fraction of SNPs in 10Kb bins. SNPs more than 100Kb away from their closest TSS are not shown for clarity. **B)** Enrichment of eQTLs and GWAS hits in promoter and enhancer elements annotated in the FANTOM project (left), and in promoter-like, TSS proximal enhancer-like, and TSS distal enhancer-like elements annotated in the ENCODE project (right). For each annotation, the enrichment value is computed as the fraction of SNPs in the annotation divided by the fraction of all SNPs in the annotation. **C)** Enrichment of eQTLs and GWAS hits in ENCODE promoter elements (left panel) and Roadmap enhancer elements (right panel) as a function of the quintile of their target gene LOEUF score (a measure of selective constraint, [63]). For promoter elements, linking to genes was done taking the closest TSS within 1Kb. For enhancers, enhancer-gene links from Liu et al. were used [53]. In panels (B) and (C), the black dashed lines mark the value of 1 on the y-axis. In all panels, values corresponding to SNPs matched for MAF, LD score, and gene density are also presented. For GWAS hits and eQTLs, error bars show 95% confidence intervals as determined by quantile bootstrapping. For matched SNPs, error bars show mean values and 95% confidence intervals in 1000 sampling iterations.

Consistent with these trends, although both GWAS hits and eQTLs are enriched in genomic regions annotated with promoter and enhancer activity by the ENCODE and FANTOM projects (Figure 5B), the relative enrichment in promoter domains versus enhancer domains is much stronger for eQTLs than GWAS hits. Also, eQTLs enrichment in both promoter and enhancer elements sharply increases as selective constraint on the target gene relaxes, particularly exceeding GWAS enrichments for the least constrained genes (Figure 5C; see Methods for details on linking regulatory regions to target genes). These results suggest that a promoter versus enhancer hypothesis (i.e., eQTLs being promoter variants, while GWAS hits being enhancer variants) does not fully explain the differences in clustering near TSSs in Figure 5A.

## 3 A model for variant discovery in GWAS and eQTL assays

In summary, we have shown that GWAS hits are systematically different from eQTL hits in important ways, including that:

- GWAS hits are enriched near selectively constrained genes (high pLI); eQTL hits are depleted.
- GWAS hits are enriched at genes with complex regulatory landscapes across different cell types, and at genes with more co-expression partners; eQTL hits are relatively depleted at all these features.
- GWAS hits are enriched across a wide variety of GO categories and among transcription factors; eQTL hits are depleted from many GO categories, and depleted among transcription factors.
- GWAS hits show only slight promoter enrichment; eQTL hits are strongly enriched at promoters.

These stark differences between GWAS hits and eQTLs may seem surprising, especially given the expectation that most noncoding GWAS hits should be eQTLs in some cell type. (Other mechanisms such as splicing do contribute, but likely in smaller proportions.) We now describe a model that predicts these qualitative patterns and shows that incomplete colocalization should be expected.

### Single cell-type model

We start by examining the case where there is only a single relevant cell type for the phenotype of interest, and eQTL mapping has been conducted in this cell type. This would be most relevant to studies where specific cell types can be studied by cell-sorting or single-cell sequencing.

For simplicity, we assume that *all* genetic effects on phenotypes are mediated via cis-effects on gene expression:

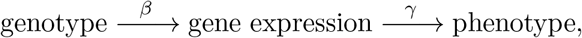

where *β* is the per-allele effect size of genotype on expression, and *γ* is the effect size of a unit change in expression on the phenotype. The net phenotypic effect of the variant on phenotype is therefore *β_γ_*.

Whether a variant is discovered as an eQTL or a GWAS hit depends on how much of the variance it explains in gene expression and in phenotype; this is given by 2*p*(1 – *p*)*β*^2^ and 2*p*(1 – *p*)*β*^2^*γ*^2^, respectively, where *p* is the allele frequency. In expectation, the variants that we discover satisfy the conditions

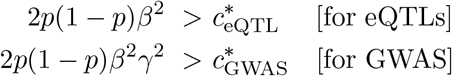

where *c*\* is a study-dependent discovery threshold. (Specifically, 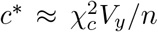, where 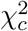 is the relevant chi-square critical-value for significance, *V_y_* is the total variance in the trait, and *n* is sample size. Equivalently, in expectation, the discovered variants satisfy the condition 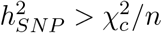, where 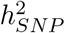 is the fraction-of-variance explained by the tested variant; see Methods.)

Consequently, the power in any given study depends on the specific parameters of the study: the fraction-of-variance explained by GWAS hits is usually much smaller than for cis-eQTLs (10^3^-fold smaller for GWAS), but this is roughly balanced by the much-larger sample sizes typical in GWAS (~10^3^-fold larger; Supplementary Note). Given these two different approaches to variant discovery, the key question is: **Should we expect GWAS and eQTL mapping to find the same hits?**

#### GWAS and eQTL mapping have power in different parts of the parameter space

We can address the question of GWAS-eQTL overlap in terms of what parts of the parameter space have appreciable power for each assay (Figure 6A). For eQTLs, we discover variants provided that 2*p*(1 – *p*)*β*^2^ is large enough; so conditional on *p* the discovery region is given by vertical contours, as shown in red. For GWAS, we need the product *β*^2^*γ*^2^ to be large enough, and so the discovery region is given by a curved blue line. Some hits are discovered by both assays because both *β*^2^ and *γ*^2^ are large enough (purple). Most importantly, some hits are discovered in GWAS only (blue region) because *β*^2^*γ*^2^ is large, but are not detected as eQTLs because their effects on expression alone (*β*^2^) are small. If we change the sample size *n* for either study, this shifts the positions of the discovery contours, but not their shapes.

**Figure 6:**
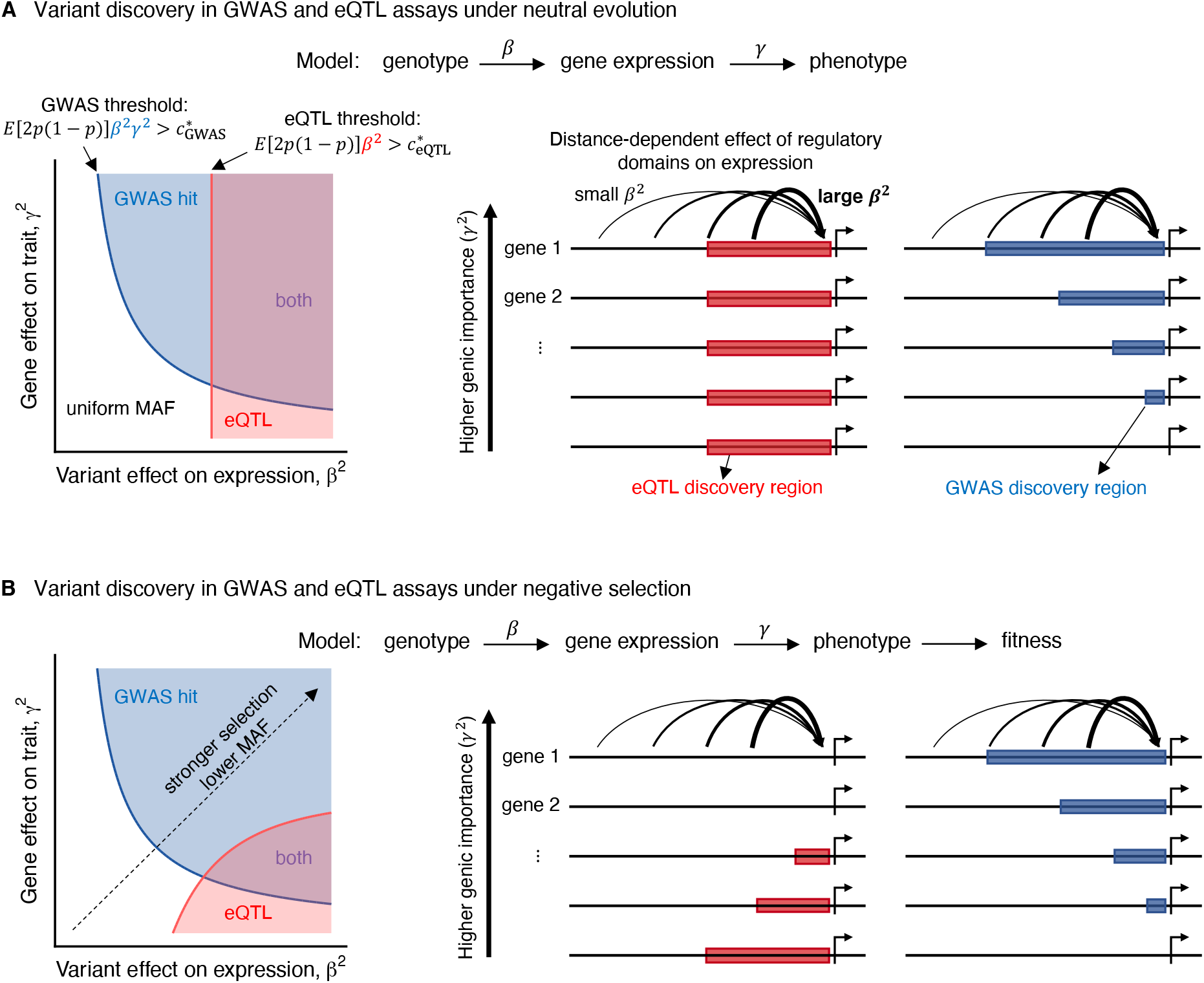
A model for variant discovery in GWAS and eQTL assays. **A)** Case of a neutrally evolving phenotype. (Left panel) Variant discovery in the space defined by variant effect on expression, β, and genic effect on phenotype, γ. Shading colors represent parameter space for the discovery of GWAS hit only (blue), eQTL only (red), and both types (purple), determined conditional on E[2p(1 – p)] which is independent of the effect sizes in the neutral case. (Right panel) schematic of variant discovery mapped to cis-regulatory domains as a function of genic contribution to the phenotype. **B)** Case of a phenotype under selection. Same as panel (A), but now the the discovery regions in the left panel are determined based on how E[2p(1 – p)] covaries with effect sizes under a model of selection (see Methods for details).

#### Selection further shrinks the region of overlap

So far we have described a neutral model, but many papers have shown selection against variants that affect complex traits [40–44]. In this scenario, selection keeps variants with large phenotypic effects at low frequencies. We derived how the discovery regions change under selection by modeling a negative correlation between *β*^2^*γ*^2^ and *E*[2*p*(1 – *p*)] (Methods; Figure 6B).

Given that selection strength is inversely related to the magnitude of the phenotypic effect, this does not systematically change the expected rankings of variants discovered in GWAS compared to the neutral scenario (Figure 6B), although it leads to a more uniform distribution of heritability across variants with intermediate and large effects (Figure S5A, a phenomenon referred to as “flattening” [40,44]). However, for eQTLs with a given magnitude of regulatory effect, *β*^2^, selection is stronger on variants acting on genes with larger *γ*^2^ values (Figure S5B). This lowers discovery power for regulatory variants linked with phenotypically important genes.

In summary, selection affects the GWAS and eQTL assays differently, mostly preventing the discovery of eQTLs at important genes. This shrinks the overlap in the discovery regions of the two assays compared to the scenario without selection (Figure 6A,B).

#### Principles of gene regulation make eQTLs more proximal

We next considered how features of gene regulation may affect GWAS-eQTL overlap. Previous eQTL studies have demonstrated that variants close to the target gene’s TSS tend to have larger effect sizes than more distal variants [61,62]. Similarly, promoter-enhancer contact frequencies decay with genomic distance between the enhancer and promoter elements [64, 65]. Thus, in terms of our model, average *β*^2^ should decay with distance from the TSS.

Under this scenario, eQTLs will be skewed towards regulatory elements near the TSS (with large *β*^2^), and depleted from distal elements (with small *β*^2^) regardless of the phenotypic importance of the target genes (*γ*^2^) (Figure 6A). GWAS hits, however, will be skewed towards phenotypically important genes (with large *γ*^2^) in a distance-dependent manner: distal regulatory elements are more likely to include a GWAS hit if acting on genes with larger phenotypic effects (Figure 6A). These patterns are predicted to be even more pronounced with selection, as large-effect TSS-proximal eQTLs should be most depleted from highly important genes, thereby further reducing eQTL– GWAS overlap (Figure 6B).

In the Supplementary Note we further discuss the role of gene regulatory architecture.

### Multi cell-type model

So far we have described the simplest case where there is only a single relevant cell type. But in practice, most current eQTL studies sample across mixtures of cell types (e.g., whole tissues), and/or environmental contexts. More fundamentally, genetic effects on a phenotype may be mediated by gene expression in multiple regulatory contexts (cell types, developmental stages, environmental stimuli, etc.).

To explore these scenarios, Figure 7 outlines a simple model of GWAS and eQTL mapping in a bulk tissue context. We derive a one-dimensional representation of this scenario in order to use the insights gained from our single-context model discussed earlier. Here, *β*_agg_ denotes an aggregate effect of a genetic variant on expression across contexts, and *γ*_agg_ denotes an aggregate effect of gene expression levels across contexts on the phenotype (Figure 7A; Supplementary Note).

**Figure 7:**
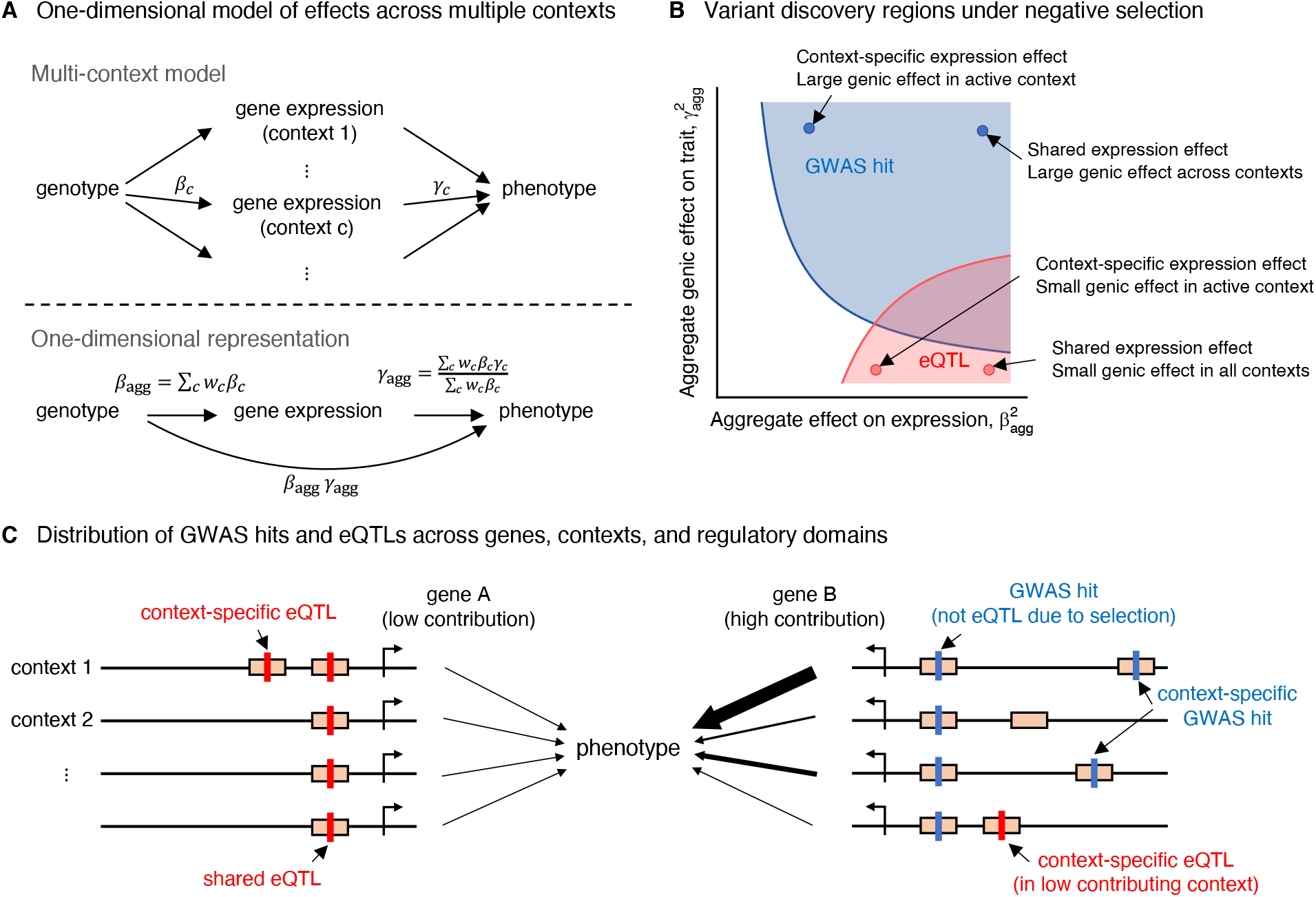
A model for variant discovery across multiple contexts. **A)** One-dimensional representation of the pathway from genotype to phenotype mediated by gene expression across multiple contexts (e.g., cell types in a tissue), by defining parameters β_agg_ and γ_agg_. **B)** Variant discovery in the space defined by βagg and γ_agg_ for a phenotype under selection; same as in Figure 6B. Shading colors represent parameter space for the discovery of GWAS hit only (blue), eQTL only (red), and both types (purple). **C)** Schematic of variant discovery across multiple contexts mapped to regulatory domains (represented by orange rectangles) for two genes at different ends of the phenotypic importance axis.

#### eQTLs are biased toward less-important genes and less-important contexts

Similar to the previous analysis, we can gain insight into the GWAS and eQTL discovery process by considering variants in terms of *β*_agg_ and *γ*_agg_ (Figure 7B). GWAS will tend to detect variants if *β*_agg_*γ*_agg_ is large enough, and this can occur through a combination of large expression effects and/or large phenotypic effects. But eQTL mapping will be most powerful for detecting large – shared – expression effects, and thus the two types of assays may have limited overlap. Figure 7C illustrates how this may play out for different variants: GWAS hits will be skewed towards functionally important genes and highly contributing contexts. Crucially, eQTLs will be skewed towards unimportant genes; meanwhile, eQTLs discovered at important genes will be skewed toward low contributing contexts. Thus, we may even find context-specific eQTLs for the “right” genes but at the “wrong” variants in the “wrong” contexts.

## 4 Discussion

Most GWAS hits are in the noncoding portion of the genome, and they are highly enriched within active chromatin. It is generally believed that the large majority of these act via effects on cis gene regulation. However, it has long been recognized that a large fraction of noncoding GWAS hits do not colocalize with known eQTLs [15, 16, 66]. This raises the question: Where are the missing eQTLs?

Certainly, part of the colocalization gap is due to the fact that we do not yet have complete measurement of all cell types, and some is due to other regulatory mechanisms such as SNP effects on splicing. But here we argue that a more fundamental issue is that GWAS and eQTL mapping are powered to identify different types of variants.

We argue that any explanation for limited colocalization must account for the fact that GWAS variants differ from eQTL variants in many important axes. Using carefully matched analyses, we have shown here that GWAS hits are biased toward more constrained genes, toward genes with functions in many GO categories, toward transcription factors, and toward genes with complex regulatory landscapes; eQTLs are biased away from all these categories. In contrast, eQTLs show a strong promoter bias that is largely absent from GWAS hits. These systematic differences cannot easily be explained by the fact that we have not yet studied all cell types (Figure S6).

Instead, to understand these observations we note that, like any mapping method, GWAS and eQTL mapping have incomplete power. In the case of eQTL mapping, a variant must explain at least a few percent of the variance to be discovered in a typical study (Supplementary Note). What types of variants, and what types of genes, are likely to cross this threshold? They tend to be variants with relatively large effects on gene regulation – large β in our model – especially in promoters. Moreover, they tend to be at genes where selective constraint is low, thereby enabling variants to drift to high frequencies.

In contrast, GWAS hits are, by design, variants that have measurable effects on an organismal trait or disease – large βγ – and hence these are biased toward functionally important genes. While selection does play an important role in reducing allele frequencies for these variants, it has a “flattening” effect, and does not systematically bias *against* discovery at important genes [40, 44].

Given these conclusions, what are the most promising routes forward for linking GWAS hits to their cognate genes and relevant cell types?

Of course, larger sample sizes in eQTL mapping will help. But for the reasons we have outlined above, the discovery regions for eQTL and GWAS mapping are systematically distinct, and extremely large eQTL samples will be needed to colocalize all desired GWAS hits (Figure S7). For less-accessible cell types this may simply not be practical. Moreover, even with extraordinarily large samples, as for blood where there is now a cis-eQTL for most expressed genes [37], our model predicts that the detected eQTLs will still be biased toward enhancers that are active in less-constrained cell types, or environmental contexts that are less relevant for interpreting GWAS.

Rather, it seems most likely that the colocalization gap will be solved with a multipronged approach, as no single method can be expected to resolve all GWAS hits. Certainly, it will be helpful to collect more cell types, more developmental stages, and larger samples. Some eQTLs may be active only in specific contexts that are underrepresented in conventional eQTL samples from bulk adult tissues such as GTEx. Likewise, some eQTLs may be represented only in rare cell types that are difficult to detect in bulk data.

Meanwhile, other types of molecular QTL assays – such as ATAC-QTLs (e.g., [67]) – may also help to plug some of the gaps. Our modeling arguments about eQTLs also hold for chromatin QTLs, but each type of assay discovers distinct variant sets, each only partially overlapping with GWAS hits and with each other (as in Figure 6B). Then, if one can identify the likely causal enhancers, these may be provisionally linked to genes using computational or functional approaches [54, 68–70].

Lastly, there are various orthogonal methods that do not suffer from quite the same discovery biases as eQTL mapping. Examples include models that predict regulatory activity of variants from DNA sequence, without the need to measure effects on gene expression (e.g., [71, 72]). Also, deciphering regulatory networks (e.g., [73–75]) can help prioritize variants that may not necessarily have large cis-effects but strong effects on the expression of tentatively functional genes. Lastly, emerging techniques using MPRA reporter assays, or CRISPR-based variant-editing or enhancer-silencing, offer powerful experimental alternatives to QTL-based techniques [76–78]. We anticipate that combinations of genome-scale techniques will ultimately help close the colocalization gap.

In summary, we have shown here that eQTLs and GWAS hits differ dramatically in several important ways. We have argued that this likely reflects essential differences in what these assays detect, shaped in large part by selection.

## 5 Methods

### Datasets

#### GWAS data

We used publicly available GWAS summary statistics for traits in the UK Biobank provided by Ben Neale’s lab (see URLs). We focused on traits with the lower bound of confidence interval on SNP heritability estimates exceeding 0.05, and with at least 50 hits (with association p-value < 5 × 10^-8^) that passed our SNP selection criteria (see section *SNP selection*). For binary traits we used heritability estimates on the liability scale, and further conditioned on traits with prevalence > 0.05. We pruned the traits list such that genetic correlation, *ρ_g_*, was < 0.5 for all trait pairs in the final list. To this end, we first sorted traits by hit count, and then starting from the trait with most hits, iterated through the list removing traits with *ρ_g_* > 0.5 with the focal trait. This procedure resulted in 44 traits listed in Table S1. Genetic correlation and SNP heritability estimates used for this procedure were downloaded from the Neale lab website.

#### eQTL data

We used eQTLs from the GTEx v8 data that were based on analyzing the subset of individuals with European ancestry in the dataset (`*EUR.signif_pairs.txt.gz` files from `GTEx_Analysis_v8_eQTL_EUR.tar` data, see URLs) to match the GWAS data. To avoid overrepresentation of brain tissues, for brain-related eQTLs we retained those identified in “Brain - Cerebellum” or “Brain - Cortex”, which have relatively distinct expression profiles among the brain regions [79]. In total 38 tissue were included listed in Table S1. To match the GWAS data, all genomic coordinates were mapped to the hg19 assembly using LiftOver [80]. In one analysis (Figure S6) we also used eQTLs detected (i) in fetal brain samples by Aygün et al. [81], (ii) at multiple stages of iPSCs differentiation towards neuronal fate by Jerber et al. [22], and (iii) in single-cell analyses of blood cell types by Yazar et al. [26]. In all analyses we conditioned on eQTLs whose target eGenes were among 18,332 protein-coding genes (see section *Gene selection*)

#### UK Biobank (UKB)

We used the UKB resource, specifically for the computation of allele frequencies, and as a linkage disequilibrium (LD) reference panel. For these purposes, we used quality control (QC) measures provided by UKB to select participants who: their reported gender, “Submitted.Gender”, matched their “Inferred.Gender” from genotypes; were not identified as heterozygosity outliers (“het.missing.outliers”== 0); did not have excessive number of relatives in the data (“excess.relatives”== 0), and were not predicted to carry sex chromosome aneuploidies (“putative.sex.chromosome.aneuploidy”==0). We further restricted our analysis to individuals identified by UKB to be of “White British” (WB) ancestry (“in.white.British.ancestry.subset”==1), and to be unrelated (“used.in.pca.calculation”==1). A total of 337,123 individuals passed these filters and were used for MAF computation. We randomly selected 10,000 of these participants as a LD reference panel.

### Gene selection

We selected 18,332 genes (Table S2) that (i) were annotated as protein-coding, and (ii) were linked with a HUGO Gene Nomenclature Committee (HGNC)-approved gene nomenclature (i.e., linked with a HGNC ID) in the GENCODE Basic gene annotations (release 39, see URLs). We used the HGNC IDs to link genic features from multiple resources to avoid issues with regard to gene names mismatching.

### SNP selection

For our SNP selection process we started with the list of 13.7 million variants that passed quality control measures for the UK Biobank analyses released by the Neale lab (labeled as “imputed-v3 Variant QC” in the Neale lab pipeline). We further applied the following filters: biallelic autosomal SNP; MAF > 0.01 among the unrelated WB individuals in the UKB (see section *Datasets*); polymorphic in the 1000 Genomes Project phase 3 data (used by Alkes Price’s lab for LD score regression, see URLs). This yielded 8,136,100 filtered SNPs.

In both GWAS and eQTL data, we first extracted SNPs that were among the list of filtered SNPs above. We then performed LD-based clumping separately for each trait in the GWAS data, and for each gene-tissue pair in the eQTL data. To this end, we used plink’s (version 1.90b6.12) --clump flag using the same parameters in both data types: p-value threshold of 5 × 10^-8^, LD threshold of *r*^2^ = 0.1 and physical distance threshold of 1Mb. The UKB resource was used as the LD reference panel (see section *Datasets*). We refer to the resulting clumped SNPs as “lead SNPs”. To make the comparisons between GWAS hits and eQTLs consistent, for both sets of lead SNPs, we removed (i) SNPs in LD (*r*^2^ > 0.8) with predicted protein-truncating or missense mutations (annotated as “ptv” or “missense” using the Variant Effect Predictor, VEP, in the Neale lab data), to condition on SNPs putatively acting through gene regulation, and (ii) SNPs > 1 Mb away from the transcription start site (TSS) of any of 18,332 protein-coding genes (see section *Gene selection*), which are not tested for eQTLs in GTEx. We further removed SNPs in the MHC region (chr6:28477797-33448354). This resulted in 22,119 GWAS hits across traits, and 118,996 eQTLs across all gene-tissue pairs (Tables S3,S4).

For both the GWAS hits and the eQTLs selected above, we included control SNPs in most of our analyses. To this end, similar to our ascertainment procedure for GWAS hits and eQTLs, we extracted 6,971,257 SNPs (from the 8,136,100 filtered SNPs) that were not in LD (r^2^ > 0.8) with predicted protein-truncating or missense mutations, were not > 1 Mb away from the TSSs of the 18,332 protein-coding genes, and were not in the MHC region. From this set, we randomly sampled 1000 SNPs for each GWAS hit or eQTL matching for MAF, LD score, and gene density (see section *SNP annotations* for the definitions and matching scheme).

### Gene annotations

We compiled a number of genic features from various resources:

- *Basic annotations*. We computed total gene length using genomic locations for transcription start and end sites extracted from GENCODE Basic annotations (see URLs). We also retrieved total coding sequence length for the longest transcript of genes from Ensembl’s BioMart tool (see URLs). These annotations were used as covariates in our logistic regression models (see section *Statistical methods*).
- *Selective constraint*. We used pLI and LOEUF measures of intolerance to LoF mutations extracted from gnomAD.v2.1’s pLoF metrics by gene data ([45, 63], see URLs).
- *Enhancer features*. We considered enhancer-gene links based on two different approaches: (i) links inferred by Liu et al. based on correlation of chromatin marks with gene expression [53] (see URLs), and (ii) links predicted based on the activity-by-contact (ABC) model from Nasser et al. [54] (see URLs). In both data, we first complied the union of all enhancer intervals per gene per biosample. We then computed two features for a given gene: (i) the number of biosamples in which the gene is linked with at least one enhancer interval (i.e., count of active biosamples), and (ii) the total length of intervals averaged across active biosamples. Genes not present in the data were assigned the value 0 for both features.
- *TSS count*. We analyzed promoter regions identified by the FANTOM consortium using Cap Analysis of Gene Expression (CAGE) [55]. We downloaded combined hg19 CAGE peaks from FANTOM5 phase 1 and phase 2 data (see URLs). For a given gene, we computed the number of peaks linked with the gene. Genes not present in the data were assigned the value 0.
- *Connectedness in co-expression networks*. We analyzed the co-expression networks from Saha et al. [56] for 16 tissues in GTEx v6p data (available from GTEx portal, see URLs). We first analyzed each tissue separately, focusing on total expression connections (“TE-TE” edges in the data) between protein-coding genes in the Transcriptome-Wide Networks (TWNs). We used the igraph package [82] in R [83] to rank genes by the number of neighbors they have in the tissue-specific networks. In the case of genes having the same number of neighbors, we used the sum of absolute weights for all edges linked with individual genes to break ties. Genes not present in a network were assigned the rank of the last gene in the network plus one. We then computed the rank-product of genes (i.e., product of the ranks of a given gene) across the 16 tissues, to construct a conntectedness measure (genes with lower rank-product have higher conntectedness).
- *Gene Ontology (GO) annotations*. We focused on 577 GO biological process (BP) terms with at least 400 genes. To this end, we first obtained specific GO annotations linked with genes using the biomaRt package [84] from the Bioconductor project (version 3.13) (attribute go_id). We then used these gene-GO links in the topGO package (version 2.44, [85]) (as the `gene2GO` parameter) to extract all genes associated with GO terms (using the `genesInTerm` function).
- *Transcription factors (TFs)*. We downloaded a list of 1639 putative human TFs from Lambert et al. [86] (see URLs).
- *Connectedness in protein-protein interaction (PPI) networks*. We used the Genoppi package [87] in R to retrieve scored InWeb PPI data (loading the inweb_table data) [88]. We then used igraph to compute the number of interactions per protein weighted by the interaction confidence scores.

To aggregate data across all resources, we first converted gene identifiers (gene symbols or Ensembl gene IDs) to HGNC IDs, and then linked all features to the selected 18,332 protein-coding genes. We used NCBI’s Gene resources (see URLs) to link HGNC IDs to gene symbols, including the official/recommended symbol as well as other used symbols labeled as “synonyms”. We used the Bioconductor’s biomaRt to link HGNC IDs to Ensembl gene IDs in the most recent Ensembl version (version 105) as well as the archived versions.

### SNP annotations

We compiled a number of SNP annotations:

- *Minor allele frequency (MAF)*. We used the UKB data (see section *Datasets*) to compute MAFs within unrelated individuals identified as White British.
- *Linkage disequilibrium (LD) score*. We used the ldsc software (see URLs, [89]) to compute LD scores using a window size of 1 centiMorgan (cM) (specified with the flag --ld-wind-cm 1). To this end, we used genetic distances (in cM) as provided by the Price lab with the “baseline (v1.1)” LD annotations for SNPs in the 1000 Genomes Project phase 3 data (see URLs). For the LD reference panel, we used 10,000 randomly selected unrelated White British individuals in the UKB (see section *Datasets*).
- *Gene density*. For a given SNP, we computed gene density as the number of protein-coding genes with their TSS falling within the 1 Mb window (±500Kb) around the SNP. The TSS coordinates were extracted from the GENCODE Basic annotations (see section *Gene annotations*).
- *Closest gene assignment*. We linked each SNP to the protein-coding gene with the closest TSS to the SNP. Subsequently, all genic features of the closest genes were assigned to the SNPs. The TSS coordinates were extracted from the GENCODE Basic annotation (see section *Gene annotations*).
- *Overlap with promoter/enhancer elements*. We considered two sets of putative promoter and enhancer elements: (i) mapped by the FANTOM5 consortium using Cap Analysis of Gene Expression (CAGE), and (ii) mapped in the phase 3 of the ENCyclopedia Of DNA Elements (ENCODE) project based on epigenetic signatures. We downloaded “permissive” enhancer regions and combined promoter regions from FANTOM5 phase 1 and phase 2 data (see URLs). In the ENCODE project, epigenetic signatures and proximity to TSSs were integrated to categorize candidate cis-regulatory elements (cCREs) as promoter-like, proximal enhancer-like (within 2Kb of nearest TSS), and distal enhancer-like (> 2Kb away of the nearest TSS) elements. We used ENCODE’s Registry V2 of cCREs (see URLs) to download the corresponding regions. Regions reported in GRCh38 were mapped to hg19 using liftOver. For our analysis in Figure 5C, we further linked the promoter-like regions to nearest TSS conditional on the distance of the regions midpoints to TSSs were < 1Kb. We constructed indicator variables for belonging to all of the above regulatory elements.

### Statistical methods

#### Estimation of mean SNP features

For a given set of SNPs (e.g., pooled set of all GWAS hits) and for a given feature (e.g., binary indicator of falling in enhancer elements) we computed mean values across all SNPs in the set. For genic features (e.g., binary indicator of belonging to a given GO annotation) we first linked SNPs to genes with the closest TSSs, and then computed mean feature values corresponding to the linked genes over all SNPs.

#### Bootstrap confidence intervals

We computed bootstrap confidence intervals for mean SNP features estimated for pooled set of GWAS hits across all traits and pooled set of eQTLs across all tissues. To construct bootstrapped samples, we first sampled traits (for GWAS hits) and tissues (for eQTLs) at random with replacement, and concatenated the sets of SNPs corresponding with sampled traits and tissues. We then sampled with replacement from the set of independent LD blocks (inferred by Berisa et al., [90]) that contain GWAS hits or eQTLs, and then concatenated the sets of SNPs (resulted from the previous bootstrapping step) belonging to the sampled LD blocks. We performed this procedure 1000 times to construct 1000 bootstrapped samples. We computed mean SNP features across all 1000 samples, and determined confidence intervals as the the range between *2.5^th^* and 97.5^th^ percentiles.

#### Control SNPs

For all GWAS hits and eQTLs, we selected control SNPs matched for MAF, LD score, and gene density. For a given SNP, we extracted SNPs (among a total of 6,971,257 SNPs and excluding the focal SNP; see section *SNP selection*) (i) with the same gene density as the focal SNP, (ii) with MAF within 0.02 of the focal SNP’s MAF, and (iii) with LD score within 0.1 standard deviation (estimated across all SNPs) of the focal SNP’s LD score. We then sampled 1000 times from this set at random with replacement to construct 1000 instances of control SNPs per SNP of interest. These were then combined across all SNPs in a set of GWAS hits or eQTLs to give 1000 sets of matched SNPs. For all gene or SNP features studied in this paper, we computed mean values across matched SNPs for all 1000 matched samples, and used the distributions to compute and report (i) mean values and confidence intervals (as the the range between *2.5^th^* and 97.5^th^ percentiles), or (ii) Z scores (see section *Analyses of individual traits and tissues*).

#### Joint analysis of features

In Figure 3B, we jointly considered the effect of multiple genic features in classifying GWAS hits or eQTLs from random SNPs. To this end, we constructed an indicator variable for GWAS hits or eQTLs (labeled 1s) versus 100,000 SNPs chosen at random from the full set of 6,971,257 SNPs (labeled 0s). We then used a logistic regression framework to predict this indicator variable using the genic features of interest. Genic feature values were normalized. We included the following covariates in the regression: MAF, LD score, gene density, absolute distance to nearest TSS, total gene length, total length of gene coding sequence, as well as dummy variables for 20 quantiles of MAF, LD score, gene density, and absolute distance to nearest TSS.

#### Analyses of individual traits and tissues

In Figures 2B and 4A we studied gene features separately for individual traits and tissues. For a given feature (e.g., proportion of SNPs that are near high pLI genes), we computed mean values for the sets of GWAS hits for individual traits and eQTLs for individual tissues. We also computed mean feature values in sets of matched SNPs (1000 sets of SNPs for each set of GWAS hits and eQTLs; see section *Control SNPs*), and used the distributions to compute (i) enrichment values as the estimated values for GWAS hits or eQTLs divided by the matched samples mean, and (ii) Z scores as the matched samples mean subtracted from the estimated values for GWAS hits or eQTLs divided by the matched samples standard deviation.

#### Analyses of eQTLs by rank

For the analysis in Figure 2C we grouped eQTLs by their rank based on association strengths (i.e., p-values) in individual tissues. To this end, in all 38 tissues, we first bin ranked eQTLs by association p-values in groups of 1000 eQTLs: first top 1000 eQTLs as group 1, second top 1000 eQTLs as group 2, and so on. We then pooled eQTLs across tissues by the ranked bins.

#### Selection of broadly unrelated GO annotations

In Figure 4A we show enrichment Z scores for GWAS hits and eQTLs across 41 GO annotations. GO annotations are hierarchical and thus interdependent. We selected these 41 gene categories by pruning 577 GO biological process annotations (see section *Gene annotations*) to give a set of broadly unrelated categories while retaining those relevant to the traits and tissues studied here. To this end, we first determined enrichment Z scores for all 577 categories (Table S6). Then, we selected the top category (i.e., most enriched or depleted) for each tissue and trait, conditioning on GO annotations with < 3000 associated genes. This gives 52 unique categories. We then pruned this set as follows: we sorted categories by the count of associated genes in an ascending order, and iterated over categories starting with the category with the least number of genes. At each iteration, we retained the focal category if gene associations with that category could not be well-predicted from the previously included categories, defined as the AUC value of < 0.75 estimated using penalized logistic regression (as implemented in the glmnet package in R using the cv.glmnet function for cross-validation, [91]) over all protein-coding genes. This gives 27 broadly unrelated categories. Using the same procedure, we built upon this set by iterating over the rest of the categories that were not among the top categories with respect to GWAS hit or eQTL enrichments, resulting in 41 categories (Table S5).

#### Analysis of non-GTEx eGenes

In Figure S6, we analyzed eGenes identified (i) in fetal brain samples by Aygün et al. [81], (ii) at multiple stages of iPSCs differentiation towards neuronal fate by Jerber et al. [22], and (iii) in single-cell analyses of blood cell types by Yazar et al. [26]. For each sample, we computed the proportion of high pLI genes among the eGenes. We then sampled the same count of genes as eGenes, 10,000 times at random from the set of all protein-coding genes. We computed the proportion of high pLI genes in each set of random genes and used the distributions to compute (i) enrichment values as the estimated values for eGenes divided by the random samples mean, and (ii) Z scores as the random samples mean subtracted from the estimated values for eGenes divided by the random samples standard deviation. For comparison, we performed the same procedure for GTEx eGenes in brain and whole blood tissues.

### Modeling variant discovery

Here we describe how we derived the discovery regions for GWAS and eQTL assays (shown in Figures 6A,B):

#### Effect of selection

A quantitative treatment of the role of selection is beyond the scope of this paper: it requires knowledge of the joint distributions of variant effects on gene expression (*β*), genic effects on phenotypes (*γ*), and allele frequency (*p*) as a function of the selection strength on the phenotype. We make simplistic assumptions to gain a *qualitative* understanding of how selection affects variant discovery in GWAS and eQTL assays.

Under a neutral model, the effect sizes and allele frequencies are independent. Thus for a given pair of (*β*, *γ*) values, the expected contribution of variants to phenotypic variance *E*[2*p*(1 – *p*)*β*^2^*γ*^2^|*β*, *γ*] = *E*[*V_p_*|*β*, *γ*]*β*^2^*γ*^2^ ∝ *β*^2^*γ*^2^ (Figure S5A), where we defined *V_p_* = 2*p*(1 – *p*) as the variance in allele frequency. Under selection, *E*[*V_p_*|*β*, *γ*] is negatively correlated with *β*^2^*γ*^2^. This is supported by empirical evidence [43, 92], as well as modeling work on the effect of selection on genetic architecture of complex traits [44]. To mimic the qualitative effect of selection, we used an asymptotic exponential form to describe the relationship *E*[*V_p_β*^2^*γ*^2^|*β*, *γ*] ∝ *κ*(1 – *e*^-*β*^2^*γ*^2^/*κ*^) (Figure S5A). In the main text we set *κ* = 2.986; under this model such a *κ* reduces *E*[*V_p_*] compared to the neutral case by ~ 10% assuming that *β* and *γ* are drawn form independent standard Normal distributions. We show in Figure S8 that as long as *E*[*V_p_*|*β*, *γ*] decreases with increasing phenotypic effect size, i.e., *β*^2^*γ*^2^, our qualitative conclusions are robust to (i) the mathematical form describing the relationship between expected phenotypic variance and effect size, and (ii) the magnitude of the relative reduction in phenotypic variance due to selection compared to neutrality.

#### Estimation of discovery regions

Given a GWAS or eQTL assay discovery threshold, *c*\* as defined in the main text, the power to discover a given variant depends on its *β*, *γ*, and *E*[*V_p_*|*β*, *γ*]. The previous section describes *E*[*V_p_*|*β*, *γ*] for a given (*β*, *γ*) pair (under selection or neutrality). Therefore, in principle, discovery regions can be solved for sets of *β* and *γ* values that satisfy 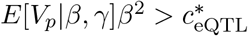 for eQTLs, and 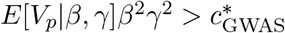 for GWAS hits.

For discovery regions in Figures 6,7, we set 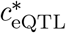 and 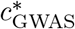 thresholds such that under the assumption that *β* and *γ* are drawn form independent standard Normal distributions, the power in GWAS and eQTL assays are matched (i.e., same fraction of causal SNPs is discovered in both assays). We set the power threshold at 15% for the visibility of discovery trends, such that there is incomplete overlap between GWAS and eQTL regions. [In Figure S7 we vary the discovery threshold to mimic the effect of sample size.] To this end, we first sampled 10 million pairs of (*β*, *γ*) ~ *N*(**0**,**I_2_**). For each pair we computed (i) the expected contribution to variance in phenotype, *E*[*V_p_*|*β*, *γ*]*β*^2^*γ*^2^, and (ii) the expected contribution to variance in expression, *E*[*V_p_*|*β*, *γ*]*β*^2^, taking *E*[*V_p_*|*β*, *γ*] values based on the exponential equation described in the previous section for the selection scenario, and equal to 1 for the neutral scenario (the ranking of variants is invariant to the scaling of *E*[*V_p_*]). For each of the four distributions (i.e., variance in phenotype and expression, in the presence or absence of selection) we computed the discovery threshold, *c*\*, as the 85^*th*^ percentile of the distribution. To plot the regions delineated by these *c*\* values, we specified a grid of *β* and *γ* values from 0 to 3.84 (corresponding to the 95^*th*^ percentile of *γ*^2^ and *β*^2^ ~ *χ*^2^(1)), with 0.0025 increments. For each pair of (*β*, *γ*) we computed expected contributions to variance in phenotype and expression as described above for the 10 million random pairs. We then identified the regions on the grid with values greater than *c*\*. The borders of the regions were smoothed using the `loess` function in `R`, applied to points on the grid between the 84.9^*th*^ and 85.1^*th*^ percentiles of the distributions of variance in phenotype or expression used to determine each *c*\*.

#### Dependence of discovery thresholds on sample size

In this section, we derive the dependence of discovery thresholds on assay sample size, *n*, and other basic parameters. We consider a simple linear regression model, estimating the effect of a single SNP, *β*, on a quantitative phenotype *Y* (a gene’s expression level or a complex trait):

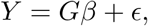

where *G* is the genotype, and 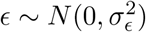 is the noise term capturing the effect of environment as well as other causal SNPs (i.e., genetic background). In OLS regression, the effect estimate 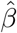, is Normally distributed with expectation 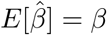. The variance of 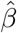 is:

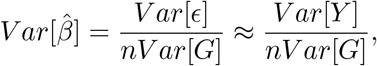

where we made the assumption that the contribution of individual SNPs to phenotypic variance is small such that *Var*[*ϵ*] ≈ *Var*[*Y*]. Now the effect is deemed significant if the *Z* score is large enough:

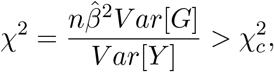

where 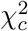 is the significance threshold. (The conventional GWAS threshold of p-value= 5 × 10^-8^ corresponds to 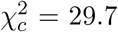). By definition, *c*\* is the critical value such that for discovered variants *Var*[*G*]*β*^2^ = 2*p*(1 – *p*)*β*^2^ > *c*\*. Now, this condition is in expectation satisfied if 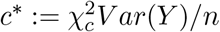.

Also, defining 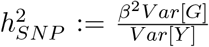 as the fraction of trait variance explained by the SNP effect, discovered variants in expectation satisfy:

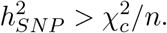

### URLs

Neale lab UKB data: http://www.nealelab.is/uk-biobank

GTEx data: https://gtexportal.org/home/datasets

NCBI’s gene_info file: ftp://ftp.ncbi.nih.gov/gene/DATA/GENE_INFO/Mammalia/Homo_sapiens.gene_info.gz

GENCODE Basic annotations: https://www.gencodegenes.org/human/release_39lift37.html

Ensembl’s BioMart: http://uswest.ensembl.org/biomart/martview

gnomAD: https://gnomad.broadinstitute.org/downloads

ABC enhancer-gene links: https://www.engreitzlab.org/resources

Liu et al.’s enhancer-gene links: https://ernstlab.biolchem.ucla.edu/roadmaplinking

FANTOM5 promoters: https://fantom.gsc.riken.jp/5/datafiles/latest/extra/CAGE_peaks

FANTOM5 enhancers: https://fantom.gsc.riken.jp/5/datafiles/latest/extra/Enhancers

Transcription factors: http://humantfs.ccbr.utoronto.ca

ldsc software: https://github.com/bulik/ldsc

LD annotations: https://alkesgroup.broadinstitute.org/LDSCORE

ENCODE cCREs: https://screen-v2.wenglab.org

## Acknowledgments

This research has been conducted using the UK Biobank resource under application Number 24983. We thank the Rivas lab at Stanford University for assistance with accessing this resource. We are grateful to Jesse Engreitz, Molly Przeworski, Guy Sella, Anshul Kundaje, Yuval Simons and members of the Pritchard lab for helpful conversations and to Jesse Engreitz, Bogdan Pasanuic, Alexis Battle, Arbel Harpak, Molly Przeworski, Guy Sella, and Wilder Wohns for valuable feedback on an earlier draft of the manuscript. This work was funded by R01HG008140, R01HG011432, and U01HG012069.

## Supplementary Material

### Supplementary Note

In this note we elaborate on a few arguments relevant to our model for variant discovery in GWAS and eQTL assays described in the main text:

#### Power considerations in GWAS and eQTL mapping

For both GWAS and eQTL mapping we consider a simple regression model, estimating the effect of a single SNP on a complex phenotype or a gene’s expression level, respectively. As detailed in the Methods section, in expectation, the discovered variants satisfy: 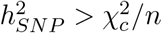, where 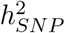 is the fraction of trait variance explained by the SNP effect, *n* is the sample size, and 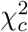 is the study-dependent discovery threshold.

The conventional GWAS threshold of p-value= 5 × 10^-8^ corresponds to 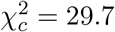. Considering a typical GWAS sample size of around 500K, discovered variants will have 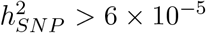. For comparison, considering a trait with a total SNP-heritability of 0.5 and 100K causal variants (which is typical of complex traits, [93]), trait variants on average have 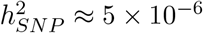.

In an eQTL study of a given gene, if we assume a p-value threshold of ~2 × 10^-4^ (on par with nominal p-value threshold values computed in GTEx at a gene-level false discovery rate threshold of 0.05), the corresponding 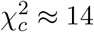. Thus, considering sample sizes on the order of 500, a variant is discovered if 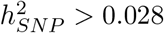. For comparison, if we assume a typical total gene expression heritability of 0.15, 30% of which is from the cis component [94] (i.e., cis-heritability of about 0.05), and 10 independent cis-regulatory variants per gene, regulatory effects on expression on average have 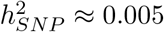. Also, if we assume the top regulatory variants per gene typically explain 25% of total heritability [94] (~83% of cis-heritability), eGenes discovered, in expectation, satisfy cis-heritability > 0.033.

#### Covariance of gene function and regulatory architecture

Figure 6 illustrates discovery regions conditional on the *β* and *γ* values. Where actual genetic variants lie in this parameter space depends on the distributions of *β* and *γ* across variants and genes. A determinant of the distribution of *β* is the variability of regulatory architecture across genes: SNPs regulating genes with more/longer enhancers likely, on average, have smaller *β*s, due to the dispersion of cis-heritability for gene expression across more regulatory variants.

On the other hand, it’s conceivable that complexity of the regulatory architecture of a gene, in part, reflects its functional importance, e.g., developmental genes are typically regulated by several enhancers elements [95]. Under this assumption, genes with more/longer enhancers, on average, have higher γs. Therefore, such genes are expected to lie in the top-left region of the parameter space (i.e., small β, large γ), and to be enriched in GWAS and depleted in eQTL assays.

These trends are similar to those predicted by Wang and Goldstein [49], though our argument does not rely on the assumption that multiple enhancer elements regulating the same gene are functionally redundant.

#### Multi cell-type model

In Figure 7 we provide a one-dimensional representation of a multi-context scenario. As described in the main text, we assign parameters *β*_agg_ denoting an aggregate effect of a genetic variant on expression across contexts, and *γ*_agg_ denoting an aggregate effect of gene expression levels across contexts on the phenotype (Figure 7A). We define *β*_agg_ ∑*_c_ w_c_β_c_*, as the weighted sum of effects over all causal contexts, where *β_c_* denotes the genetic effect on expression in context *c* with the weighting *w_c_*. As an example, considering a situation where a gene’s expression in a tissue influences a phenotype, *β_c_* represents the effect in cell type *c*, *w* the relative abundance of different cell types, and *β*_agg_ the effect estimate in a bulk assay of the causal tissue.

We define 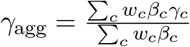, where *γ_c_* denotes the effect of a unit change in target gene’s expression in context c on the phenotype. The rationale for this definition is such that, similar to our single-context model, the quantity *β*_agg_*γ*_agg_ gives the net effect of the genetic variant on phenotype. For a given variant, the interpretation of *γ*_agg_ is the mean genic effect across contexts weighted by the effect of the variant on gene expression. For example, if a variant is active in a single-context *c*, *γ*_agg_ is *γ_c_*, and if the variant has the same effect on expression across all contexts *γ*_agg_ is ∑*_c_ w_c_γ_c_*.

In this multi-context model, for a given variant, the role of the degree of sharing of the expression effects across multiple contexts is analogous to the role of distance to TSS in the single-context model. In this view, using the result that discovered eQTLs are expected to be more TSS-proximal than GWAS hits, eQTLs are expected to be more shared across contexts, or on the flip side, GWAS hits are expected to be more context-specific than eQTLs.

## Supplementary Figures

**Figure S1:**
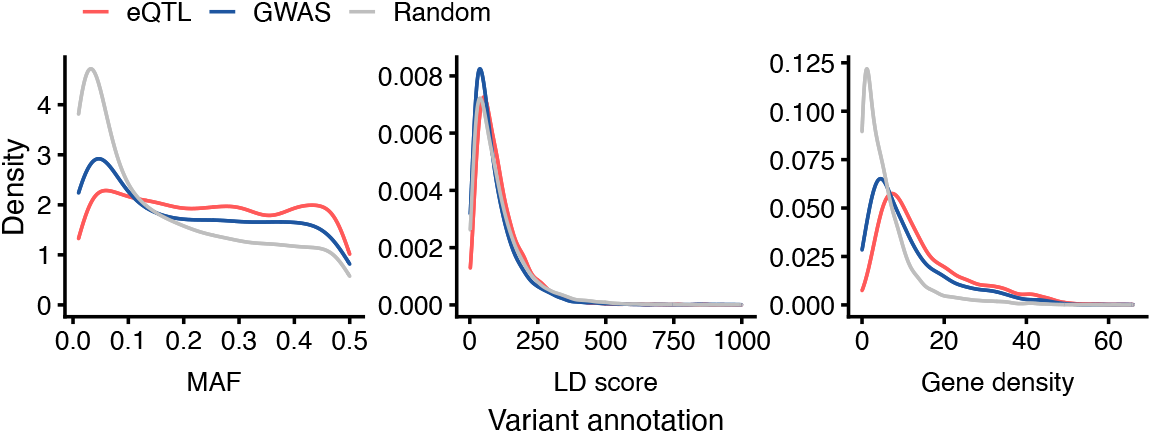
Basic variant-level differences between GWAS hits and eQTLs. Distribution of minor allele frequency (MAF), linkage disequilibrium (LD) score and gene density for eQTLs (red), GWAS hits (blue), and 100,000 randomly chosen variants. LD score values are cut at 1000 for clarity.

**Figure S2:**
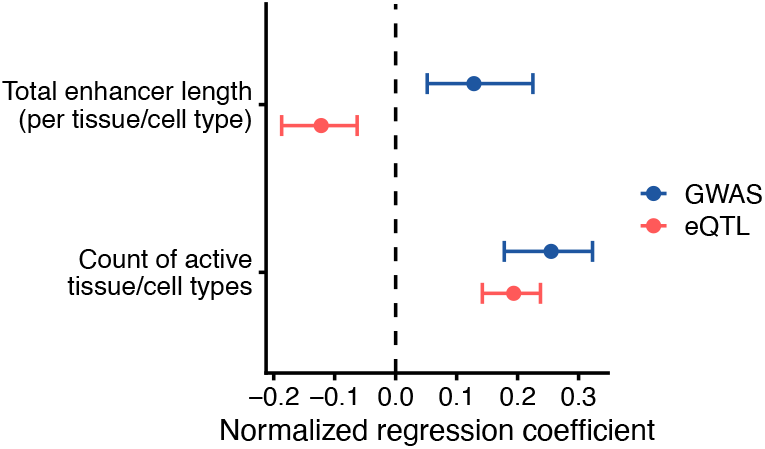
GWAS and eQTL genes have different enhancer architectures. Same as Figure 3B, but using enhancer-gene links predicted by the activity-by-contact (ABC) model from Nasser et al. [54] (Methods). For a given gene, we computed (i) the number of biosamples in which a gene has an enhancer, and (ii) the average total enhancer length (in base pairs) across active biosamples. Shown are logistic regression coefficients corresponding with the two enhancer features for predicting GWAS hits (blue) and eQTLs (red) versus random variants after adjusting for confounders (Methods). Error bars show 95% confidence intervals as determined by quantile bootstrapping.

**Figure S3:**
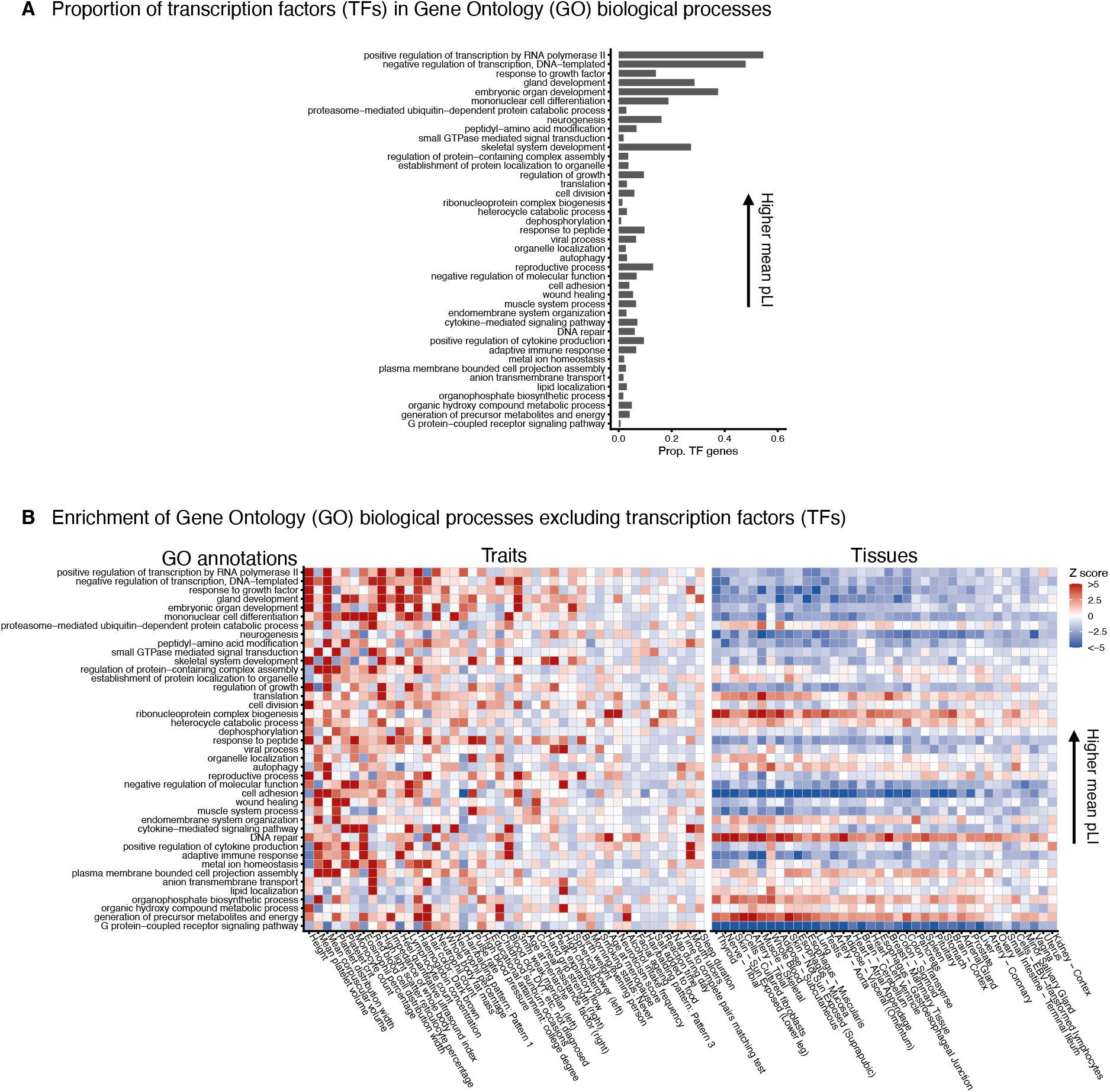
Contribution of transcription factors (TFs) in Gene Ontology (GO) annotations and their enrichment in GWAS and eQTL genes. **A)** Proportion of TFs in 41 GO biological processes shown in Figure 4A. **B)** Same as Figure 4A, but now excluding TFs from all 41 gene categories before computing enrichment values among GWAS and eQTL genes. Traits and tissues (x-axis) are sorted by hit count (decreasing from left to right), and GO annotations (y-axis) are sorted by the mean pLI value of genes within categories (before removing TFs, replicating the ordering in Figure 4A). For each trait- or tissue-GO category pair we computed enrichment Z scores based on 1000 sampling iterations of variants matched for MAF, LD score, and gene density (see Methods). The color map represents enrichment (red) or depletion (blue) of a given gene set among GWAS or eQTL genes.

**Figure S4:**
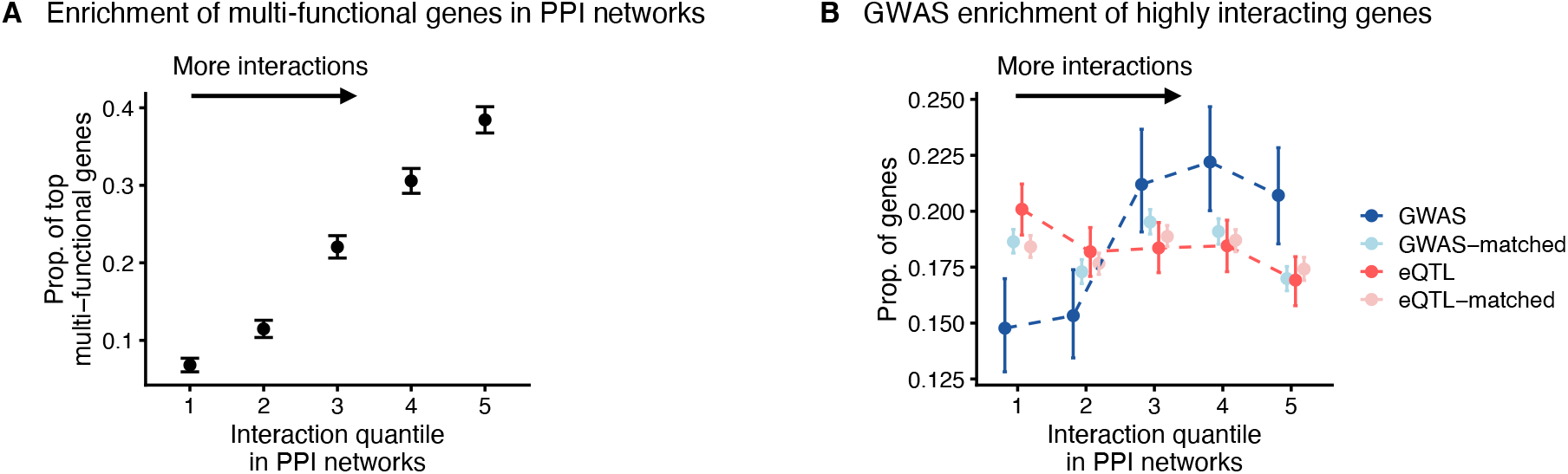
Multi-functionality of highly interacting genes in protein-protein interaction (PPI) networks and their enrichment in GWAS genes. **A)** Proportion of genes in bins ranked by the number of interactions in the InWeb PPI network [88] that are among the top multi-functional genes (defined as top 20% of genes ranked by the count of Gene Ontology (GO) categories they belong to, see Methods). Error bars show 2 standard errors. **B)** Fraction of GWAS and eQTL genes in gene bins ranked by the number of interactions in the InWeb PPI network. For GWAS hits and eQTLs, error bars show 95% confidence intervals as determined by quantile bootstrapping. For matched variants (for MAF, LD score and gene density, shown in light blue and red colors), points and error bars show mean values and 95% confidence intervals in 1000 sampling iterations.

**Figure S5:**
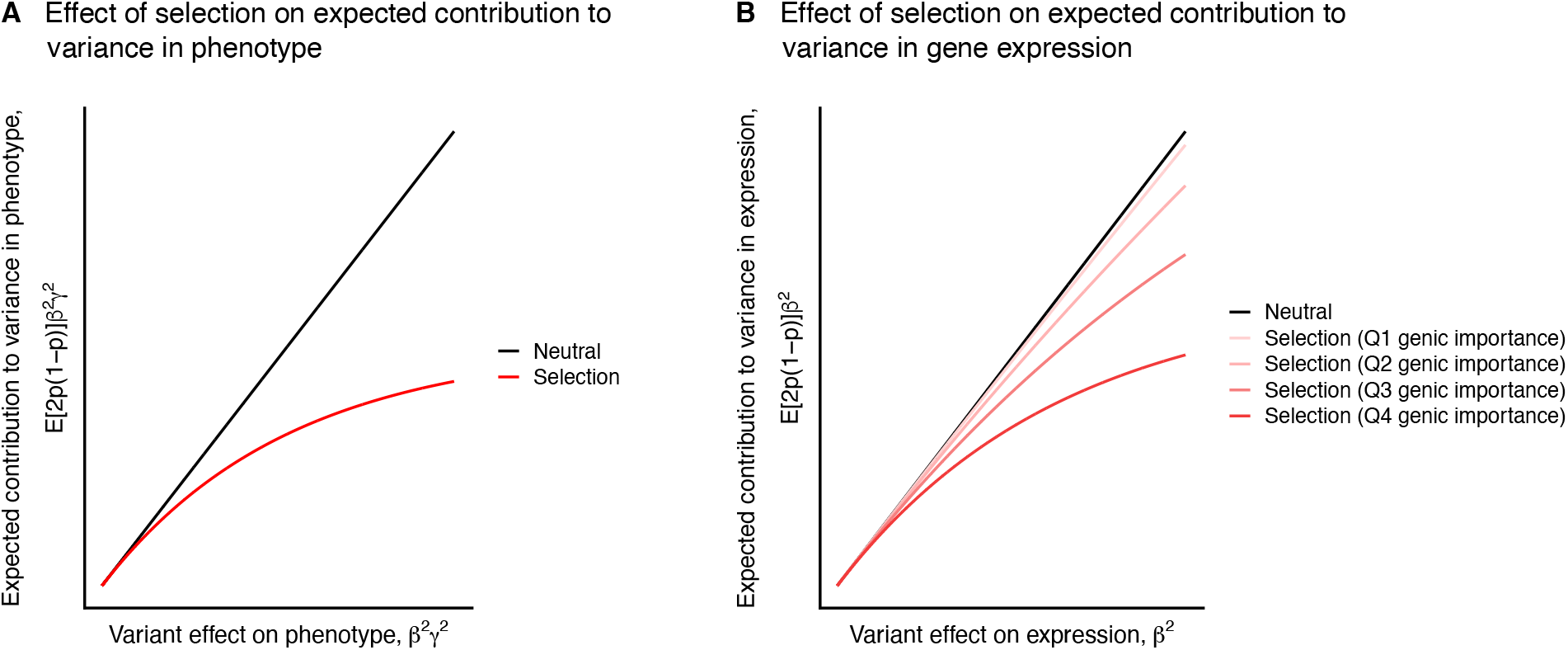
Effect of selection on variants contribution to variance in phenotype and gene expression. **A,B)** As described in the main text, we consider a model of phenotypic effects mediated by effects on gene expression intermediates: a genetic variant affects the expression of the target gene with effect β, and the gene expression intermediate affects the downstream phenotype with effect size γ. **A)** Contribution to phenotypic variance. Under a neutral model, contribution to phenotypic variance, E[2p(1 – p)]β^2^γ^2^, is proportional to phenotypic effect, β^2^γ^2^, as effect size and allele frequency are uncoupled. Selection keeps higher effect variants at lower frequencies (i.e., lowering E[p(1 – p)]) and thus “flattens” the expected contribution to variance. The red line shows a flattened curve taking E[2p(1 – p)β^2^γ^2^ |β, γ] ~ κ(1 – e^-β^2^γ^2^/κ^), with κ = 2.986 (Methods). **B)** Contribution to variance in gene expression. Similar to the argument in (A), under neutrality, contribution to variance in gene expression, E[2p(1 – p)]β^2^, is proportional to the effect on expression, β^2^. Under selection, flattening (i.e., lowering of E[p(1 – p)]) is more pronounced for variants regulating high-effect (i.e., high γ^2^) genes. Red lines show trends for four quantiles of γ^2^, where γ ~ N (0, 1); darker colors show higher γ^2^ values. See Methods for modeling details.

**Figure S6:**
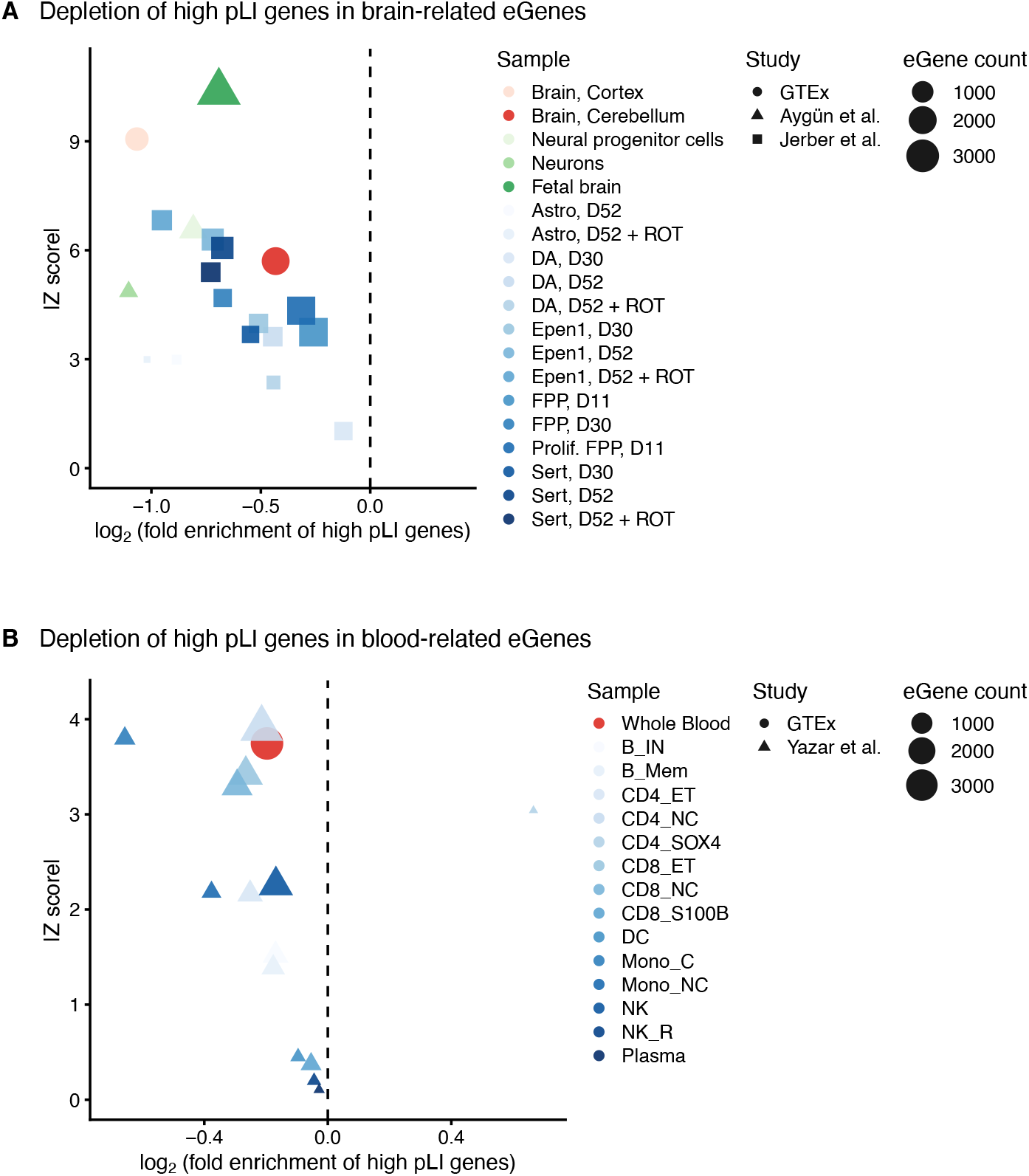
Depletion of selectively constrained genes among non-GTEx eGenes. The factors we described against the discovery of trait-eQTLs likely bias eQTL assays in any context. As proof of concept, we show that similar to GTEx eGenes, eGenes identified in non-conventional eQTL assays are also depleted of strongly selected genes. **A)** Enrichment of high pLI genes in eGenes identified (i) in fetal brain samples by Aygün et al. [81], (ii) at multiple stages of iPSCs differentiation towards neuronal fate by Jerber et al. [22], and (iii) in GTEx brain tissues. Sample labels for Jerber et al. refer to different ascertained cell types, at different days of differentiation, and in the presence or absence of stimulation by rotenone (ROT) (see Jerber et al. for details). **B)** Enrichment of high pLI genes in eGenes identified in (i) single-cell analyses of blood cell types by Yazar et al. [26], and (ii) GTEx whole blood. Sample labels for Yazar et al. refer to different blood cell types (see Yazar et al. for details). Enrichment values (on the x-axis) and Z scores (on the y-axis) were computed based on values observed in 10,000 sampling iterations of random genes (Methods).

**Figure S7:**
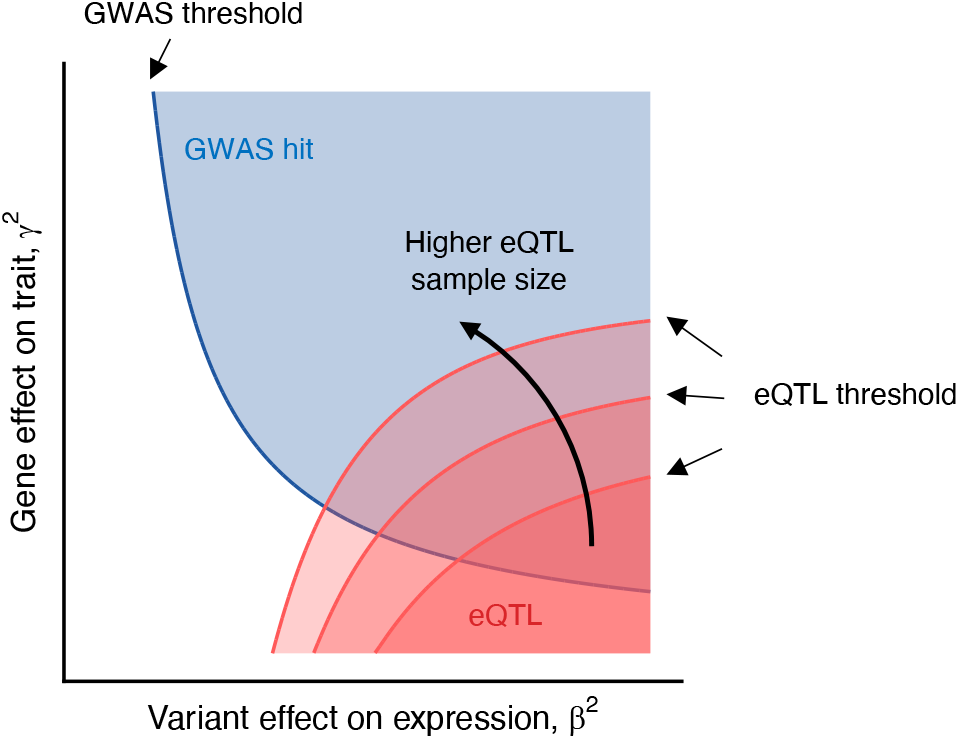
Effect of eQTL assay sample size on discovery. Same as Figure 6B, but with three eQTL discovery thresholds corresponding to different sample sizes. The discovery thresholds are derived by setting the power rate to 15% for GWAS under the assumptions detailed in the Methods section, and to 10%, 15% and 20% for eQTLs.

**Figure S8:**
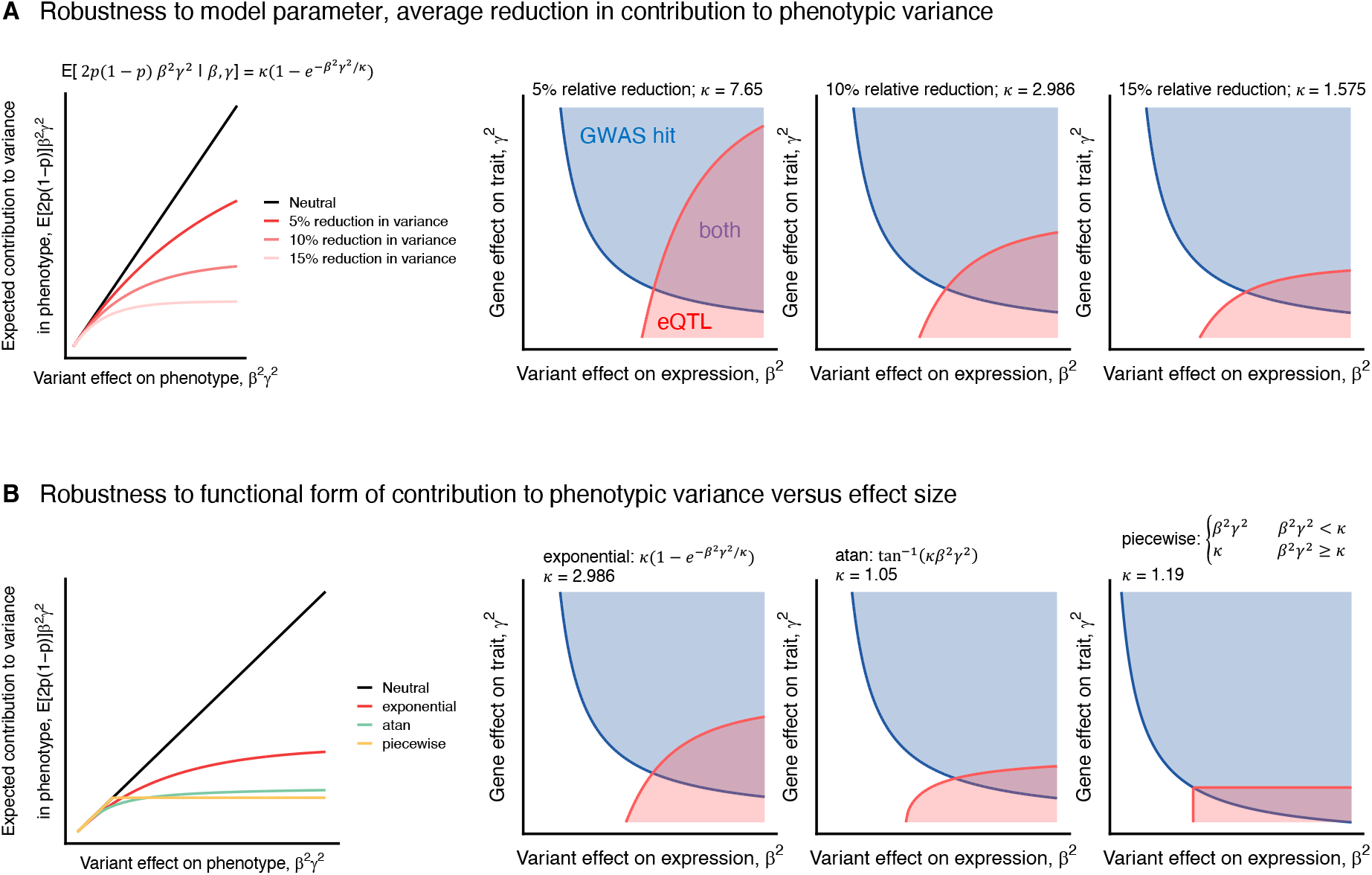
Robustness of qualitative results to modeling choices. In Figure 6B we show GWAS and eQTL discovery regions under the scenario that the phenotype is under selection. We incorporate selection by modeling a reduction in contribution to phenotypic variance (compared to the neutral scenario) as a function of the phenotypic effect (see Methods for details). For plots shown in the the main text we chose the exponential form E[2p(1 – p)β^2^γ^2^|β, γ] ~ κ(1 – e^-β^2^γ^2^/κ^), with κ = 2.986 to obtain an average reduction of ~ 10% across randomly sampled β and γ values. We show here that our qualitative results are robust to the choice of (A) average reduction in contribution to phenotypic variance and (B) the functional form of the variance under selection. In panel (B), κ values are tuned to give an average reduction in contribution to phenotypic variance of ~ 10% for all three models.

## Supplementary Tables

Table S1: **List of traits and tissues.**

Table S2: **List of autosomal protein-coding genes.**

Table S3: **List of GWAS hits.**

Table S4: **List of eQTLs.**

Table S5: **List of broadly unrelated Gene Ontology (GO) biological process terms.**

Table S6: **Enrichment of Gene Ontology (GO) biological processes in GWAS and eQTL genes for individual traits and tissues.**

